# Rational design paving the way for improving glucose tolerance and catalytic properties of a β-glucosidase from *Acetivibrio thermocellus*

**DOI:** 10.1101/2024.08.05.606605

**Authors:** Chinmay Kamale, Abhishek Rauniyar, Prasenjit Bhaumik

## Abstract

Cellulases are an ensemble of enzymes that hydrolyse cellulose chains to fermentable glucose, hence, are widely used in bioethanol production. The last enzyme of the cellulose degradation pathway - β-glucosidase, is inhibited by its product – glucose. The product inhibition by glucose hinders cellulose hydrolysis limiting the saccharification during bioethanol production. Therefore, engineered β-glucosidases with improved glucose tolerance along with the catalytic efficiency are the need of the hour. This study focuses on the rational engineering of β-glucosidase from *Acetivibrio thermocellus* (WT-AtGH1). Recombinant WT-AtGH1 exhibited activity on cellobiose and p-nitrophenyl-β-D-glucosidase as substrates and retained around 80% of its activity over 48 hours at 55°C, pH 5.5. However, WT-AtGH1 showed low glucose tolerance of 380 mM as compared to the required *IC*_50_ value of > 800 mM for industrial use. Therefore, the rational design approach was applied for improving the glucose tolerance of this enzyme. We determined 3 Å resolution crystal structure of WT-AtGH1. The structure-based engineered G168W-AtGH1 and S242W-AtGH1 mutants exhibited improved glucose tolerance of 840 mM and 612 mM, respectively. Surprisingly, S242L-AtGH1 mutant showed ∼ 2.5-fold increase in the catalytic efficiency as compared to WT-AtGH1. A combinatorial effect of improved glucose tolerance, as well as enhanced catalytic efficiency, was observed for the G168W-S242L-AtGH1 mutant. All the mutants with enhanced properties showed considerable stability at industrial operating conditions of 55°C and pH 5.5. Thus, we present the next-generation mutants of WT-AtGH1 with improved glucose tolerance and kinetic properties that have the potential to increase the efficiency of the saccharification process for second generation bioethanol production.

## Introduction

The enormous population increase worldwide has led to increased demand for energy resources – mainly conventional fuels. The conventional fuels used today as a significant energy source include gasoline, diesel and other petroleum-based fuels. The increased consumption of conventional energy resources is contributing to the rise of carbon dioxide load in the environment. Therefore, it is essential to search for alternate sources of fuel that are eco-friendly. With the population surge, there is a growing need for the management of biological waste generated through various human-related activities. Processes are being developed for sustainable management of such waste by utilizing it for the production of biofuels [1]. Bioethanol is the preferred fuel over gasoline because of its high-octane number, high efficiency of combustion, no toxicity of combustion products as well as reduced greenhouse gas emissions [2–6]. Biological wastes generated from food crops, agricultural industries and daily household activities are used as raw materials for bioethanol production. These wastes are majorly composed of lignocellulosic mass. Lignocellulosic biomass utilized as raw material for bioethanol production mainly consists of three essential components – cellulose (40-50%), lignin (15-20%) and hemicelluloses (25–35%) [7], out of which cellulose is recalcitrant and resistant to degradation because of extensive intra-fibres and inter-fibre hydrogen bonding [8].

Cellulases (EC 3.2.1.x) are enzymes belonging to the family - glycosyl hydrolase that catalyse the hydrolysis of cellulose. Cellulase is ensemble of different proteins and enzymes forming a cellulosomal assembly in many anaerobic bacteria for effective cellulose degradation [9]. The cellulose degradation pathway involves three prime enzymes – endoglucanase (EC 3.2.1.4), exoglucanase (EC 3.2.1.91) and β-glucosidase (EC 3.2.1.21) which act synergistically on cellulose to hydrolyse it completely to monomeric β-D-glucose [10]. β-glucosidase is the last enzyme of cellulose degradation pathway which converts di/tri/tetra-saccharide units generated from cellulose by the action of endo- and exo-glucanases into monomeric β-D-glucose molecules [11,12]. High concentrations of β-D-glucose have an inhibitory effect on β-glucosidase, leading to the accumulation of cellobiose and other higher oligosaccharides; which in turn slows down exo- and endo-glucanase activity. Hence, β-glucosidase acts as a control point for the cellulose degradation pathway. Glucose-mediated inhibition of β-glucosidase is critical as this affects the whole process of cellulose breakdown [13]. A study describing the process for the production of soluble sugars from biomass mentioned that the saccharification process could reach the glucose levels of 50 – 150 g/L of broth (0.55 M – 0.83 M of glucose) in the reactor [14]. Such high levels of glucose in the reactor at the industrial level pose a potential hurdle in the saccharification process as product inhibition of β-glucosidase by glucose becomes the rate-limiting step. Hence, engineering of known β-glucosidases to improve their glucose tolerance is an inevitable need for economical and efficient industrial saccharification processes for bioethanol production [15,16]. At the industrial scale, bioethanol production is carried out by two major processes – simultaneous saccharification and fermentation (SSF) and Separate hydrolysis and fermentation (SHF). In SSF, saccharification by hydrolytic enzymes and fermentation by microbes is carried out in the same reactor. In such cases, there is a disparity in maintaining the temperature and pH of the reactor as optimum conditions for cellulolytic enzymes (temperature of 50 °C and pH 5 – 6) are different from the ethanol-producing yeast (temperature of 30 – 37 °C and pH 4 – 5). This ultimately leads to low ethanol yields [17–19]. The SHF process has the upper hand over SSF as the saccharification and fermentation take place in separate reactors. This aids in carrying out the two steps at their optimum conditions, leading to high ethanol yields [20,21]. Therefore, SHF is the preferred industrial process for bioethanol production. Generally, the saccharification in SHF is carried out in the temperature range of 50 - 60 °C with pH maintained at 4.8 – 5.5 [22]. As the saccharification process runs over a period of 48 hours, cellulolytic enzymes should retain their activities at the temperature of 50 - 60 °C with pH 4.8 – 5.5 [22, 23]. Considering these limitations, the development of β-glucosidases having tolerance to high glucose concentration with stability at the industrial operating conditions of saccharification is highly essential for industrial use.

Most of the naturally occurring β-glucosidases do not possess all the above-mentioned desired properties. Hence, engineering of β-glucosidases stands out as a potential solution using two well-established strategies: (a) directed evolution and (b) rational design approach. Both strategies have been shown to be successful towards the improvement of β-glucosidases for their enhanced thermal stability as well as glucose tolerance [24–30].

This study focuses on a β-glucosidase from the thermophilic anaerobic cellulolytic bacterium – *Acetivibrio thermocellus* [31]. β-glucosidase from *Acetivibrio thermocellus* (WT-AtGH1) was cloned and expressed as well as structurally and biochemically characterized. X-ray crystallographic studies showed WT-AtGH1 has the structural characteristics of a typical glycosyl hydrolase 1 family member. Sequence and structural alignment of this β-glucosidase (AtGH1) with other glucose tolerant β-glucosidases helped in identifying the possible residues for mutations that would increase the glucose tolerance of AtGH1. The point mutations - G168W and S242W introduced in AtGH1 by the rational design approach of protein engineering led to a considerable increase in glucose tolerance of AtGH1. Additionally, S242L point mutation led to improving the catalytic efficiency of the enzyme towards cellobiose. Our structural, biochemical, kinetic and thermal stability data indicate that the generated mutants can be used for the saccharification process of bioethanol production. This study highlights the importance of the rational design approach for the protein engineering of industrially important β-glucosidase.

## Results

### Wild type (WT)-AtGH1 is a thermotolerant enzyme with low glucose tolerance

WT-AtGH1 (EC 3.2.1.21) containing the C-terminal 6X His tag (theoretical MW of the fusion protein is ∼52 kDa) was expressed in *E. coli*. A considerable amount of expression of the recombinant protein in soluble fraction was observed with induction by 1 mM IPTG at 30 °C for 18 hours. WT-AtGH1 was purified with Ni-NTA affinity and subsequently by gel filtration chromatography (**Fig. S1**).

Purified WT-AtGH1 was catalytically active, and the optimum temperature and pH for the activity were determined using pNPG and cellobiose as substrates. WT-AtGH1 showed maximum activity at 60 °C with both substrates. The enzyme showed ∼85% and ∼95% retention of the activity at 55 °C for pNPG and cellobiose, respectively, compared to the activities at 60 °C (**Fig. 1A**). Interestingly, WT-AtGH1 showed different pH optima with two substrates. With pNPG, WT-AtGH1 showed maximum activity at pH 6.5. However, the enzyme exhibited optimum activity in the range of pH 5 - 5.5 with cellobiose as a substrate (**Fig. 1B**).

**Fig. 1.**
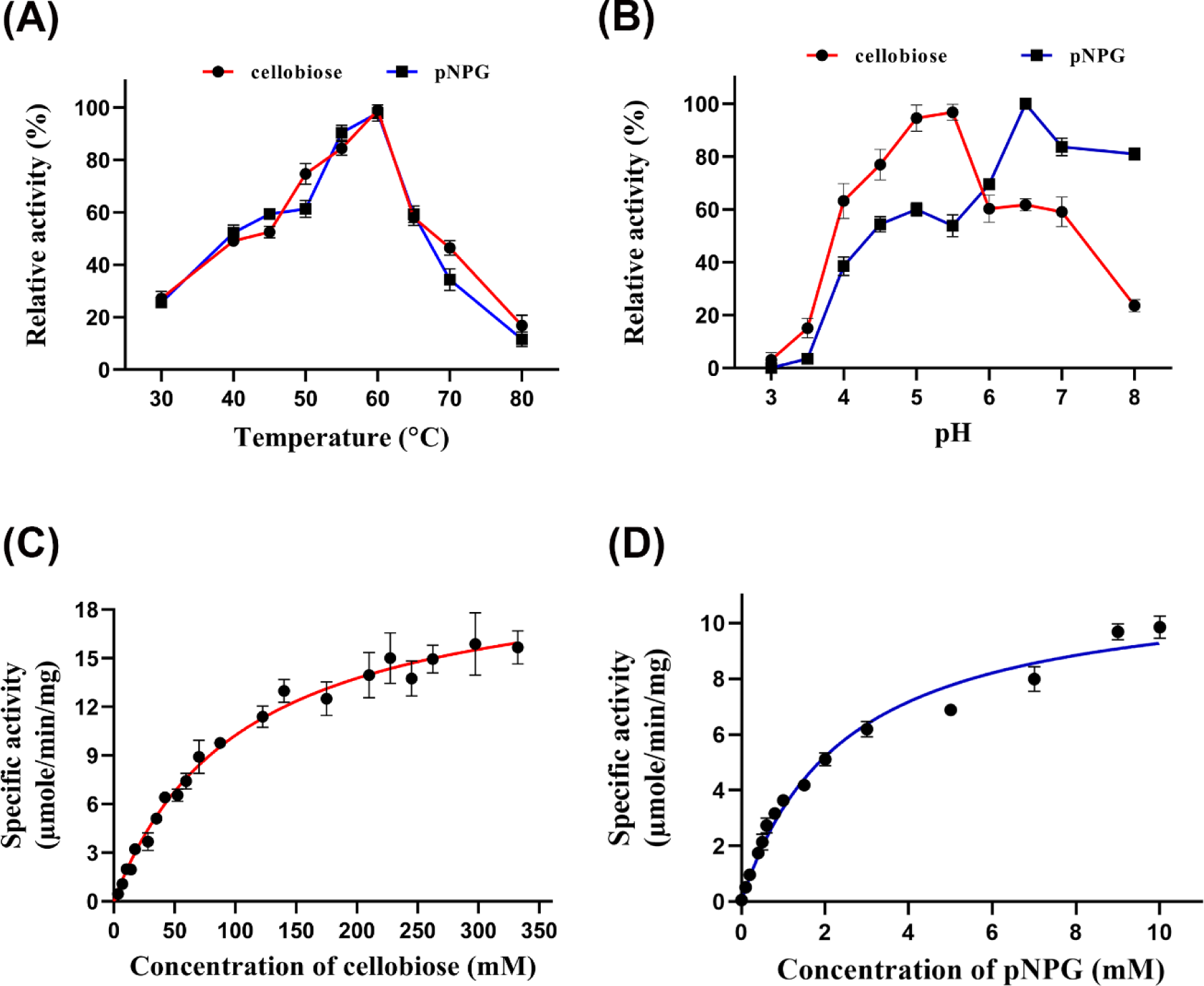
Biochemical and kinetic characterizations of WT-AtGH1. (A) Effect of temperature and (B) effect of pH on the activity of WT-AtGH1. Relative activity of WT-AtGH1 is plotted against varying temperature and pH range, respectively. Assays were done with pNPG (blue curve) and cellobiose (red curve). Kinetic characterizations of WT-AtGH1 with (C) cellobiose and (D) with pNPG. Data was fitted to the Michaelis-Menten kinetic model. All experiments were performed in triplicates (n = 3) with error bar representing ± SEM.

Kinetic parameters were determined with pNPG and cellobiose as substrates to assess the catalytic properties of WT-AtGH1. All the kinetic parameters for WT-AtGH1 (**Fig. 1C and 1D**) determined by fitting the data to the Michaelis Menten curve are summarised in **Table 1**. The substrate affinity (*K*_m_) for pNPG was 2.42 mM, lower than that of cellobiose (*K*_m_ = 105 mM). The turnover numbers (*k*_cat_) determined for pNPG and cellobiose were 604.8 min^−1^ and 1101.4 min^−1^, respectively. The catalytic efficiency, *k*_cat_/*K*_m_ with cellobiose was 10.49 mM^−1^ s^−1^ which was lower than that of pNPG. The *IC*_50_ value for the WT-AtGH1 with glucose was determined to be 380 mM (**Fig. 2A**). *IC*_50_ value is referred to as “glucose tolerance” in this study. The inhibition pattern of glucose on the activity of WT-AtGH1 was elucidated by determining enzyme activities with increasing concentrations of pNPG at the fixed concentration of glucose. The inhibition analysis by Dixon plot (**Fig. 2B**) showed glucose to be the competitive inhibitor of the enzyme with *K*_i_ of 137 mM of glucose.

**Fig. 2.**
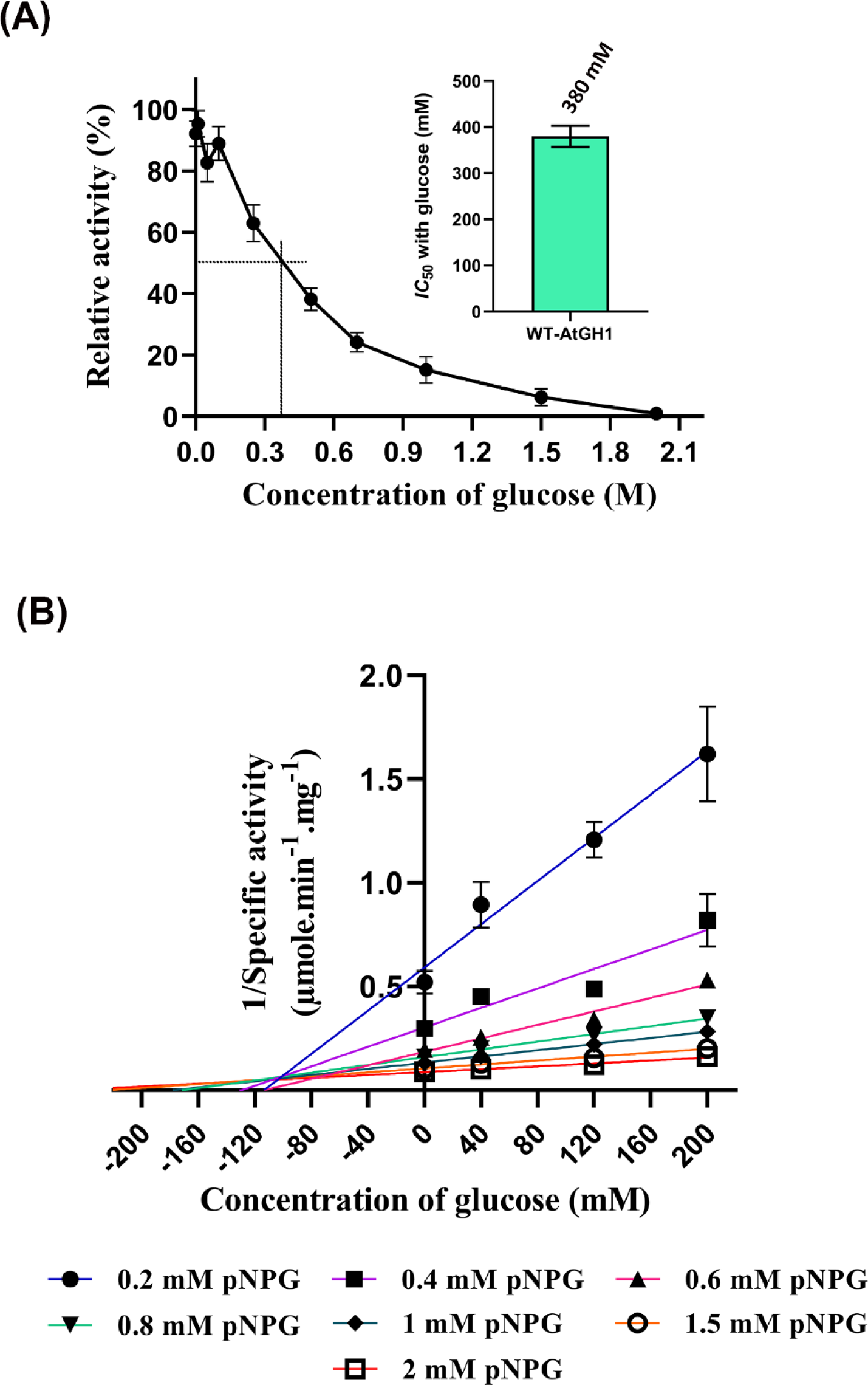
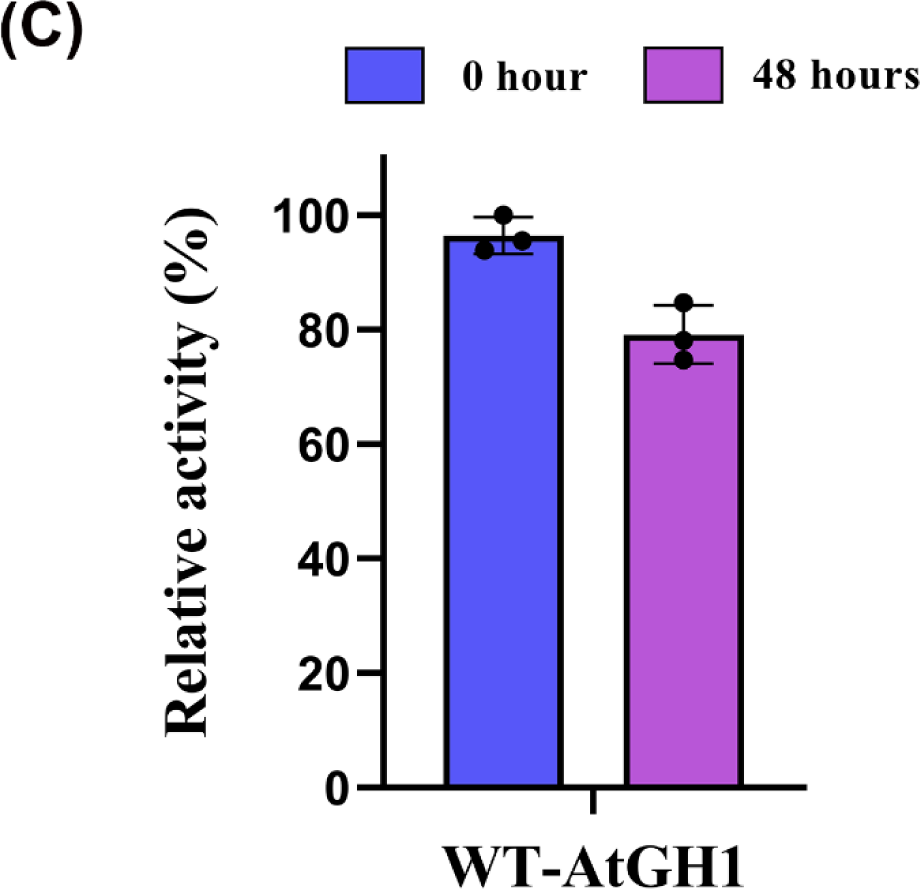
Glucose tolerance and thermal stability characterization of WT-AtGH1. (A) Effect of glucose on the activity of WT-AtGH1, graph represents the relative change in the activity of WT-AtGH1 with varying concentrations of glucose. The inset shows the *IC*_50_ value of glucose for WT-AtGH1. (B) Dixon plot for competitive inhibition of WT-AtGH1 with glucose. Assays were done with varying concentrations of pNPG as a substrate with three different concentrations of glucose (40 mM, 120 mM, 200 mM). (C) Comparison of thermal stability of WT-AtGH1 at 55 °C and pH 5.5. The bar diagram represents the relative activity of WT-AtGH1 at 0 hours (blue bar) and after 48 hours incubation (purple bar). All experiments were performed in triplicates (n = 3) with error bar representing ± SEM.

**Table 1.** Kinetic parameters of WT-AtGH1 and its mutants measured with pNPG and cellobiose as substrates (ND – Not determined)

|  | WT-AtGH1 |  | G168W-AtGH1 |  | S242W-AtGH1 |  | G168W-S242W-AtGH1 |  | S242L-AtGH1 |  | G168W-S242L-AtGH1 |  |
| --- | --- | --- | --- | --- | --- | --- | --- | --- | --- | --- | --- | --- |
|  | cellobiose | pNPG | cellobiose | pNPG | cellobiose | pNPG | cellobiose | pNPG | cellobiose | pNPG | cellobiose | pNPG |
| $K_m$ (mM) | 105 ±<br>7.73 | 2.42 ±<br>0.17 | 114.1 ±<br>6.75 | 2.23 ±<br>0.13 | 125.2 ±<br>13.3 | 2.1 ±<br>0.1 | ND | ND | 78.74 ±<br>4.67 | 1.79 ±<br>0.11 | 112.9 ±<br>5.52 | 1.32 ±<br>0.11 |
| $V_{max}$ (μmol<br>min<sup>-1</sup> mg<sup>-1</sup>) | 20.98 ±<br>0.62 | 11.52 ±<br>0.32 | 12.73 ±<br>0.31 | 10.42 ±<br>0.22 | 31.44 ±<br>1.44 | 30.99 ±<br>0.47 | ND | ND | 40.47 ±<br>0.86 | 8.1 ±<br>0.17 | 35.63 ±<br>0.72 | 14.15 ±<br>0.33 |
| $k_{cat}$ (min<sup>-1</sup>) | 1101.4 | 604.8 | 668.3 | 547.05 | 1650.6 | 1626.9 | - | - | 2124.7 | 425.25 | 1870.6 | 742.9 |
| $k_{cat}/K_m$<br>(min<sup>-1</sup> mM<sup>-1</sup>) | 10.49 | 249.9 | 5.85 | 245.31 | 13.18 | 774.71 | - | - | 26.98 | 237.56 | 16.57 | 562.8 |

Lignocellulosic biomass is highly heterogeneous and contains monovalent, divalent and trivalent metal ions along with lignin, hemicellulose and cellulose. Along with this, the source of lignocellulose, reactors used for pretreatment of biomass and water used contribute towards the presence of metal ions [32]. Hence, deciphering the effect of the commonly found metal ions on the activity of WT-AtGH1 is crucial for its industrial applicability. The WT-AtGH1 activity has been evaluated with pNPG as a substrate in the presence of selected metal ions (**Fig. S2**). Notably, Mg^2+^ and Co^2+^ showed ∼ 70 – 80% enhancement of the enzymatic activities at 1 mM concentration. However, higher concentrations (> 1 mM) of Mg^2+^ and Co^2+^ did not have any effect on the activity of WT-AtGH1. For Ca^2+^ and Mn^2+^, an increase in the concentration of metal ions led to an increase in the enzyme activity. The presence of Cu^2+^ and Hg^2+^ inhibited the activity of the enzyme to a considerable extent.

The thermal stability of WT-AtGH1 was assessed at industrial operation conditions of the saccharification reactor. With cellobiose as a substrate, WT-AtGH1 without any additive retained 71% of its activity over the incubation of 48 hours at 55°C and pH 5.5 compared to its activity at 0 hour (**Fig. 2C**). The retention in the enzyme activity of WT-AtGH1 under the above-mentioned temperature and pH conditions is indicative of the intact tertiary structure of the enzyme (**Fig. S3**).

### Structural features of WT-AtGH1

The crystal structure of WT-AtGH1 was determined with X-ray diffraction data extending to 3.0 Å resolution. Initial refinements of structure with *REFMAC5* followed by final refinement cycles with the automated refinement module of *Phenix* improved the final *R*_factor_ and *R*_free_ values. The data collection and refinement statistics are mentioned in **Table 2**. A monomer of WT-AtGH1 is the functional entity showing a characteristic structural fold of GH1 family β-glucosidases. The WT-AtGH1 shows a typical (α/β)_8_ barrel structure consisting of closely packed alternate α - helical and β - sheeted secondary structures linked together by interconnecting loops (**Fig. 3A**). The superposition of WT-AtGH1 with a β-glucosidase from *C. cellulovorans* (PDB ID: 3AHX) shows the well-conserved GH1 family fold-typical (α/β)_8_ barrel structure. The active site pocket is situated in this barrel fold (**Fig. 3B**). Two catalytic residues, Glu355 and Glu166 are situated at the end of β7 (ENG motif) and at end of β4 (NEP motif), respectively (**Fig. 3C, S5**). Out of three major active site topologies known for GH superfamilies, WT-AtGH1 shows the presence of a characteristic crater. The depth of the crater is observed to be 16 Å and at the bottom lies the two catalytic glutamate residues (E166 and E355) (**Fig. 4B**). These two catalytic glutamates are surrounded by polar residues – Asn222, His121, His180 and Gln20. The active site crater for WT-AtGH1 (**Fig. 4B**) is shown to be lined with aromatic as well as hydrophobic amino acid residues such as Tyr296, Trp328, Trp 402, Tyr412, Ile 179, Met326, and Val161. Out of these, Tyr296 was found to be conserved in all GH1 family β-glucosidases. The exact role of this tyrosine residue is still unknown. A distinct density of Tris molecule is observed in the active site pocket of WT-AtGH1. The presence of a Tris molecule at the active site can be corroborated with the use of 50 mM Tris buffer (with 400 mM NaCl) for the purification of WT-AtGH1. This Tris molecule is well-stabilized by the hydrogen bonding interactions with Glu166, Glu355 and Glu409. The stabilization of Tris at the active site of AtGH1 is further supported by the temperature B factor of 22.71 Å^2^ shown by the Tris molecule (**Fig. 3D**). The presence of Tris in the active site of GH1 family β-glucosidases has been reported in published crystal structures. Due to the presence of three hydroxyl groups, tris may be considered as substrate mimic [33–36]. The distance between two catalytic glutamate (Glu166 and Glu355) residues is ∼5.5 Å (**Fig. 4A**), suggesting that WT-AtGH1 follows the retention-type of catalytic mechanism. Along with the distance of ∼5.5 Å between two catalytic glutamates, distinct densities of water molecules are also observed. Both these water molecules – Wat1 and Wat2, are in hydrogen bonding interactions with catalytic Glu 166 and Glu355 (**Fig. S4**). Wat1 and Wat2 are closely associated with catalytic glutamate residues with temperature B factors of 5.27 Å^2^ and 8.34 Å^2^, respectively. These temperature B factors indicate that Wat1 and Wat2 are well stabilized at the active site of WT-AtGH1. Out of Wat1 and Wat2, one might be the catalytic water molecule involved in the donation of the hydroxyl group to the product during the reaction. The structure superimposition (**Fig. 4C**) of the active site crater of WT-AtGH1 with the active site crater of glucose-bound UnBGl1, a β-glucosidase from soil metagenome indicates the probable position of glucose molecule in the active site crater of WT-AtGH1 [37,38]. A hydroxyl group of anomeric carbon C1 can be seen to be in close proximity to Glu355 of WT-AtGH1. This indicated that Glu355 from WT-AtGH1 may form a covalent intermediate with one of the β-D-glucose moieties from cellobiose and, hence, may act as a base (nucleophile), and Glu166 may act as the acid in general acid-base catalysis.

**Fig. 3.**
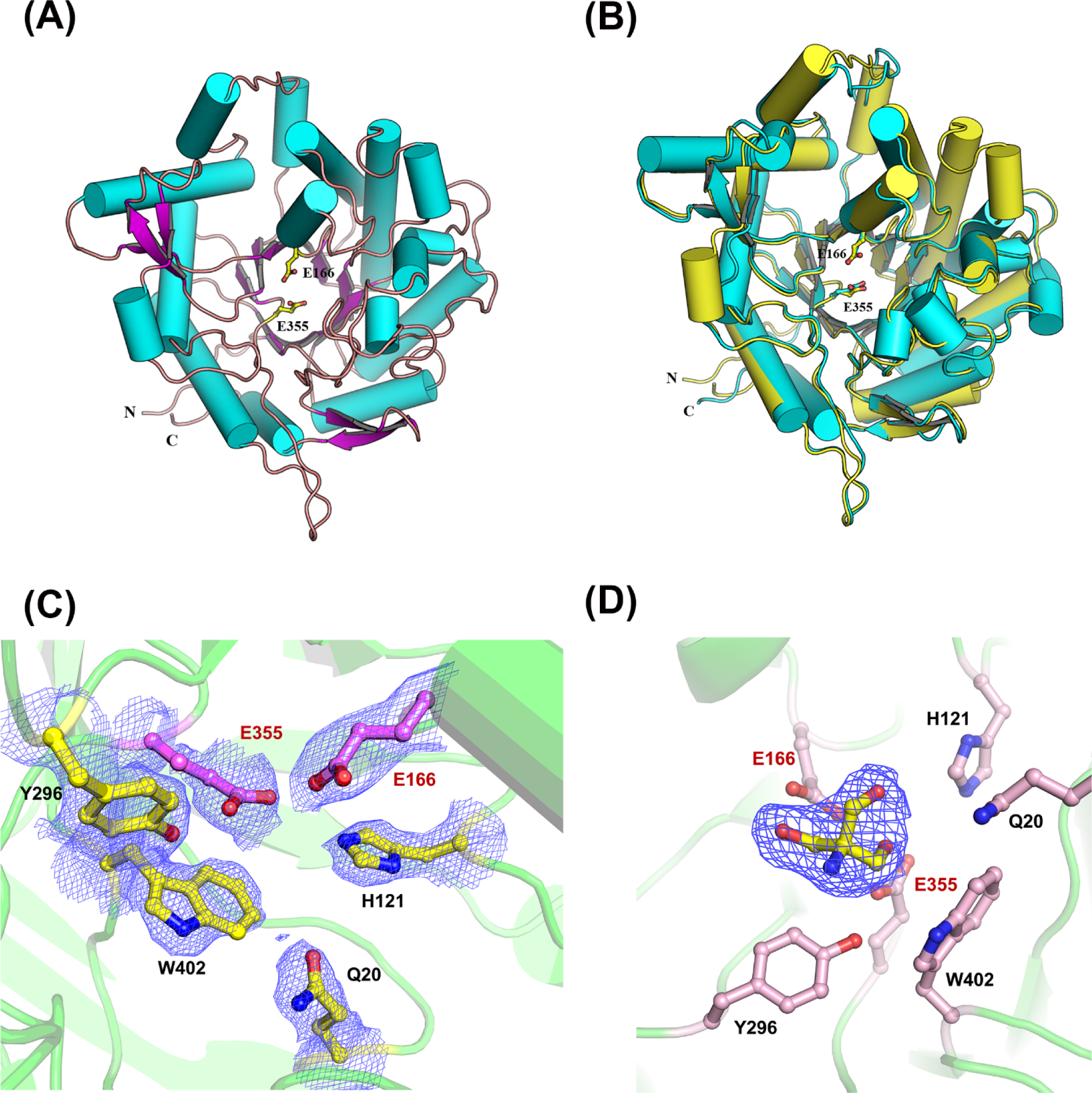
Tertiary structure and active site of WT-AtGH1. (A) Crystal structure of WT-AtGH1 monomer with typical characteristic (α/β)_8_ fold of GH1 family proteins is represented as cartoon. The α - helices shown as cylinders (in cyan) the β sheets are shown as arrows (in glucosidase from *C. cellulovorans* (PDB ID: 3AHX, cartoon representation in cyan). Both the enzymes show (α/β)_8_ fold – characteristic feature of GH1 family glucosidases, α-helices are represented as cylinders and β-sheets as arrows with catalytic glutamates in ball and stick representation. (C) 2*F*_o_-*F*_c_ map (contour level 1σ) for side chains (ball and stick) of the active site residues forming bottom of active site crater is shown in blue mesh. Two catalytic glutamates are represented in pink colour. (D) Tris molecule observed in the active site of WT-AtGH1 represented as ball and sticks in yellow surrounded by 2*F*_o_-*F*_c_ electron density map (contour level 1σ). Surrounding active site residues on WT-AtGH1 are shown as ball and stick in salmon colour.

**Fig. 4.**
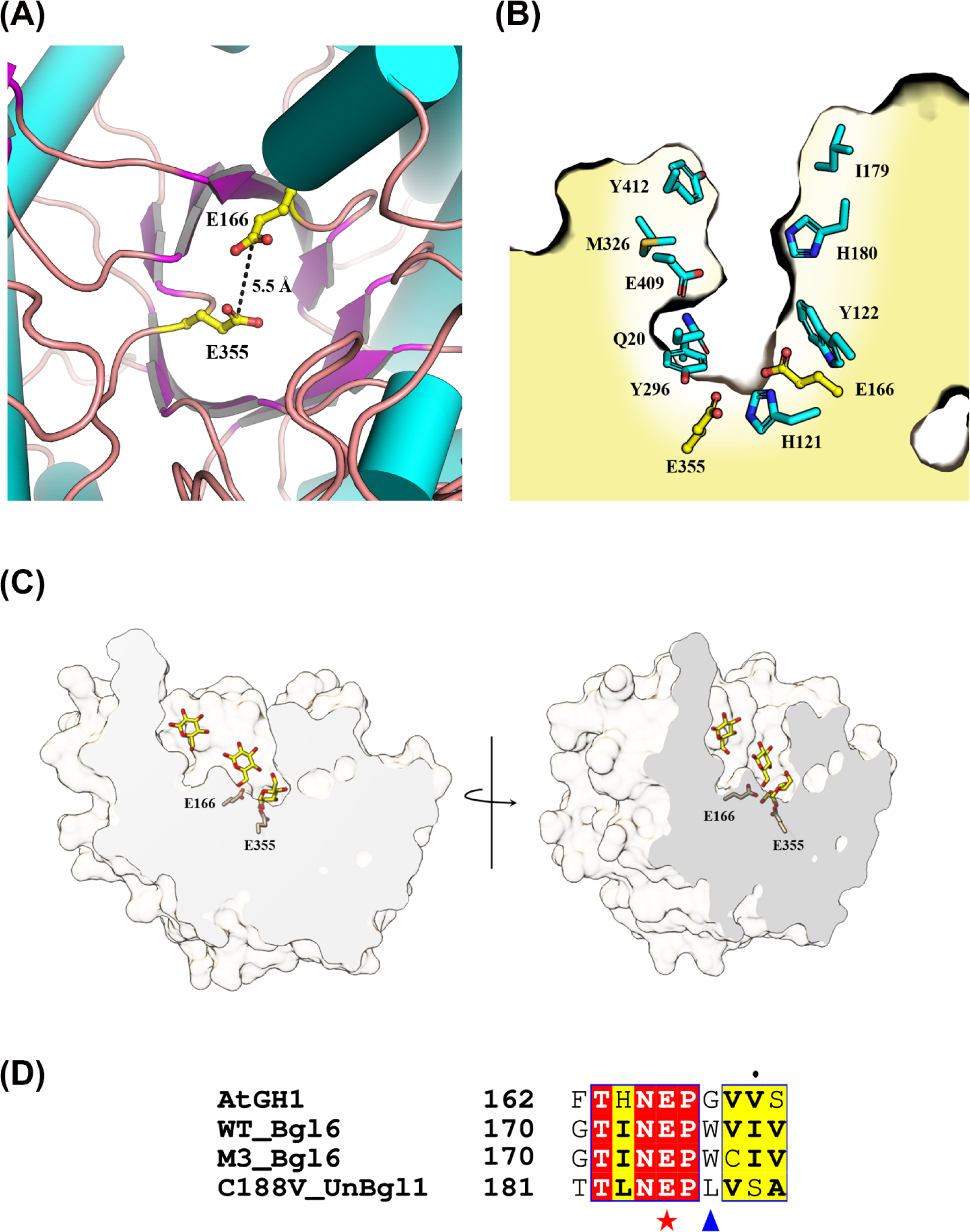
Molecular details of the active site crater of WT-AtGH1 and its comparison with other β-glucosidases. (A) Distance between two catalytic glutamates E166 and E355(ball and stick) is shown as dotted line (B) Representation of the active site crater of WT-AtGH1. Residues lining the crater are represented as sticks and the catalytic glutamates are shown in ball and stick representation. (C) Solid section of the active site crater of WT-AtGH1 with three glucose molecules (yellow carbon) bound at the active site crater of UnBGl1 structure is shown. Catalytic glutamates (light pink carbon) of WT-AtGH1 are represented as stick models. (D) Multiple sequence alignment for some of the known glucose tolerant β-glucosidases with WT-AtGH1. Red star indicates the position of one of the catalytic glutamates from NEP motif (corresponding positions of E166 from WT-AtGH1) with blue triangle marking the position of mutation for G168W of WT-AtGH1.

**Table 2.** Data collection and refinement statistics for WT-AtGH1 structure.

| **Data collection statistics<sup>a</sup>** | WT-AtGH1 |
| --- | --- |
| Space group | *P*1 |
| Unit cell dimensions<br>*a*, *b*, *c* (Å)<br>$\alpha$, $\beta$, $\gamma$ (°) | 109.93, 158.62, 158.91<br>60.12, 88.19, 87.48 |
| Resolution (Å) | 35.0 – 3.0 (3.2 – 3.0) |
| Wavelength (Å) | 1.5418 |
| Temperature (K) | 100 |
| Observed reflections | 349165 (61898) |
| Unique reflections | 175099 (31024) |
| Completeness (%) | 93.9 (94.7) |
| $R_{\text{merge}}^b$ (%) | 16.5 (64.8) |
| Mean $I/\sigma(I)$ | 4.3 (1.3) |
| $CC_{1/2}$ (%) | 97.2 (50.3) |
| Redundancy | 1.99 (1.99) |
| **Refinement statistics** |  |
| Resolution (Å) | 35.0 – 3.0 |
| Working set: no. of reflections | 166166 |
| $R_{\text{factor}}$ (%) | 30.37 |
| Test set: no. of reflections | 8748 |
| $R_{\text{free}}$ (%) | 34.16 |
| *Number of atoms* |  |
| Proteins | 44061 |
| Water | 355 |
| Tris | 16 |
| *Average isotropic B-factor ($\text{\AA}^2$) of all atoms* | 45.95 |
| R.M.S.D. |  |
| Bond length (Å) | 0.006 |
| Bond angle (°) | 1.434 |
| *Protein geometry<sup>c</sup>* |  |
| Ramchandran plot allowed (%) | 98 |
| Ramchandran plot generously allowed (%) | 1.5 |
| Ramchandran plot outlier (%) | 0.5 |
| PDB ID | 9IQB |

<sup>a</sup>Values in parentheses indicate statistics in highest resolution shell.
<sup>b</sup> $R_{\text{merge}} = \sum |I - \langle I \rangle| / \sum I$ where $I$ is the observed integrated intensity, $\langle I \rangle$ is the average integrated intensity obtained from multiple measurements, and the summation is overall observed reflections.
<sup>c</sup>Values are obtained from PROCHECK analysis.

### Structure based rational design of WT-AtGH1 for improving its glucose tolerance

Sequence and structure based comparative analyses of WT-AtGH1 with three high glucose tolerant GH1 family β-glucosidases – Bgl6, M3 mutant of Bgl6 and C188V mutant of UnBGl1 were carried out [37,38]. Multiple sequence alignment of Bgl6 and M3-Bgl6 (with *IC*_50_ of 3.5 M and 3 M of glucose, respectively), C188V mutant of UnBGl1 (*IC*_50_ of 1.8 M of glucose) with WT-AtGH1 showed conserved TXNEP motif and ENGXAX motifs, each containing one catalytic glutamate residue (**Fig. S5**). An immediate amino acid position followed by TXNEP motif shows the presence of aromatic residue – tryptophan in both Bgl6 and M3 mutant of Bgl6, whereas for C188V-UnBGl1, it showed leucine at that position (**Fig. 4D**). The structural alignment of WT-AtGH1 with Bgl6 showed that amino acid at this position was surrounded by a patch of hydrophobic residues (**Fig. 5B**). This site, known as the +1 subsite, is one of the three subsites in the active site crater of the enzyme, which is observed to bind glucose molecule at very high concentrations of glucose [37,38]. Hence, we expected an increase in the glucose tolerance of the enzyme by mutating the Gly168 at this position to Trp. Purified G168W-AtGH1 showed optimum activity in the range of pH 5.5 - 6 and in the temperature range of 55°C - 60°C (**Fig. S1 and S6**). The kinetic parameters of G168W-AtGH1 were assessed with pNPG and cellobiose as substrates. G168W-AtGH1 was able to hydrolyse pNPG and cellobiose indicating mutation did not lead to loss of activity of the enzyme. G168W-AtGH1 showed retention in the kinetic properties - *K*_m_ and *V*_max_ (with both pNPG and cellobiose as substrates) compared to WT-AtGH1 (**Fig. S7**). However, the turnover number and, hence, the catalytic efficiency of G168W-AtGH1 were lower than that of WT-AtGH1 with cellobiose as a substrate (**Table 1)**. With pNPG as a substrate, G168W-AtGH1 showed the *IC*_50_ value of 820 mM glucose. A single point mutation of G168W led to a more than two-fold increase in the glucose tolerance level of the mutant enzyme, compared to WT-AtGH1 with *IC*_50_ of 380 mM of glucose (**Fig. 6A and 6B**).

**Fig. 5.**
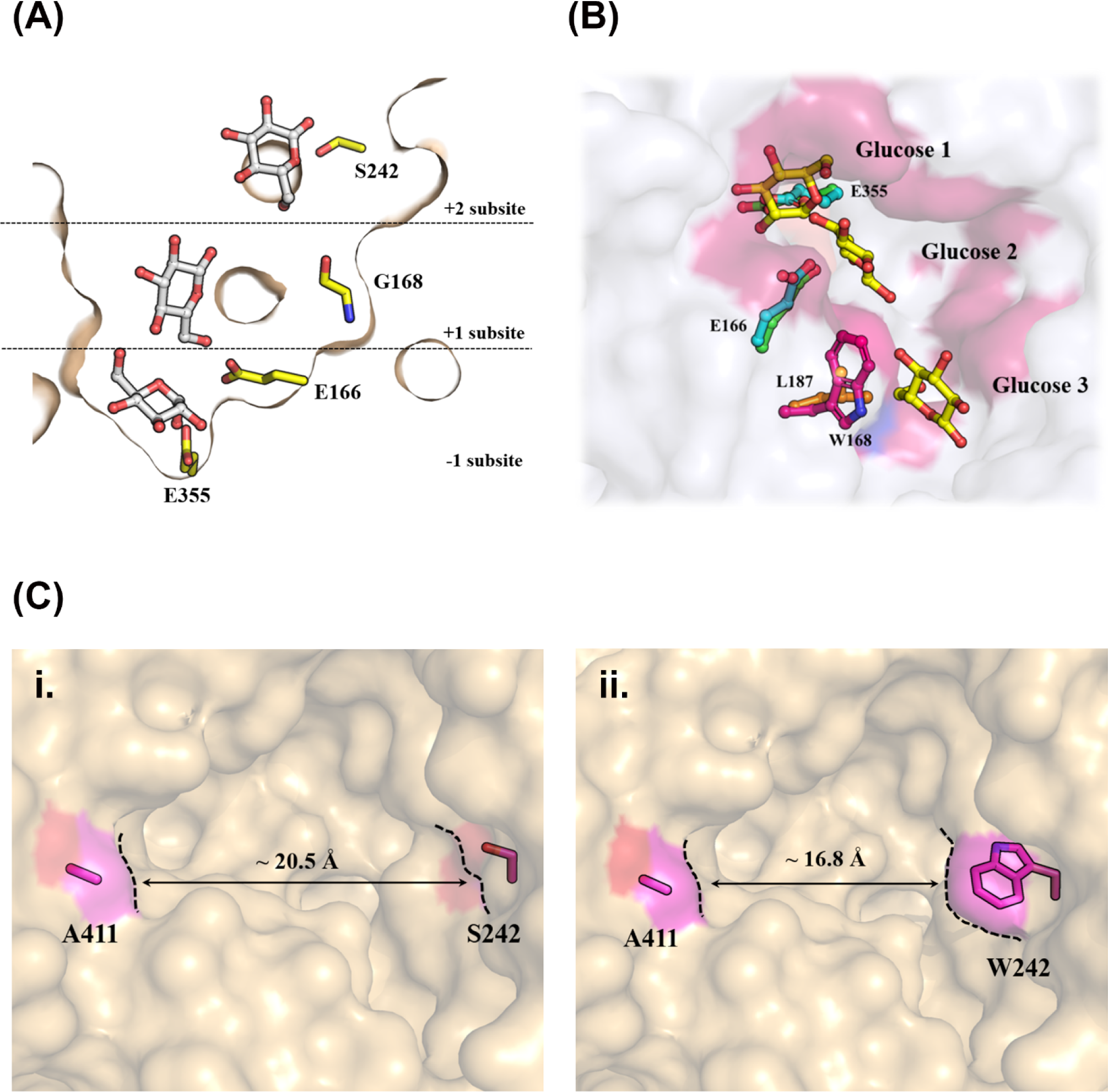
Structure based strategy for the improvement of glucose tolerance of WT-AtGH1. (A) Cross section of active site crater of WT-AtGH1. Structural superposition of WT-AtGH1 with that of glucose bound wild type UnBGl1 defined the −1, +1, and +2 subsites in WT-AtGH1. −1 site possesses E166 and E355 catalytic residues of WT-AtGH1. Glycine168 forms three subsites are occupied by glucose molecules presented as represented as ball and sticks in grey. (B) Structural superposition of WT-AtGH1 and glucose bound UnBGl1, three glucose molecules (represented as ball and sticks in yellow) bound to three glucose binding subsites in the active site crater of UnBGl1. E166 and E355 are the catalytic residues of WT-AtGH1 (represented as ball and sticks in blue) superposed with catalytic residues of UnBGl1 – E185 and E370 (represented as ball and sticks in green), respectively. G168 is mutated to bulky W168 (represented as ball and sticks in pink) manually and is shown to be superposed with the L187 (represented as ball and sticks in orange) of UnBGl1. Pink patches shown are the hydrophobic residues lining the active site crater of WT-AtGH1. (C) S242, one of the gatekeeper residues of WT-AtGH1, was proposed to keep the active site crater entry wide, which may lead to inhibition of the enzyme at low glucose concentrations. **i.** The diameter of the active site crater entry for WT-AtGH1 is ∼ 20.5 Å**. ii.** When S242 was mutated to W242, it led to a narrowing of the active site crater entry. The resultant diameter of the entry was hence determined to be ∼ 16.8 Å.

**Fig. 6.**
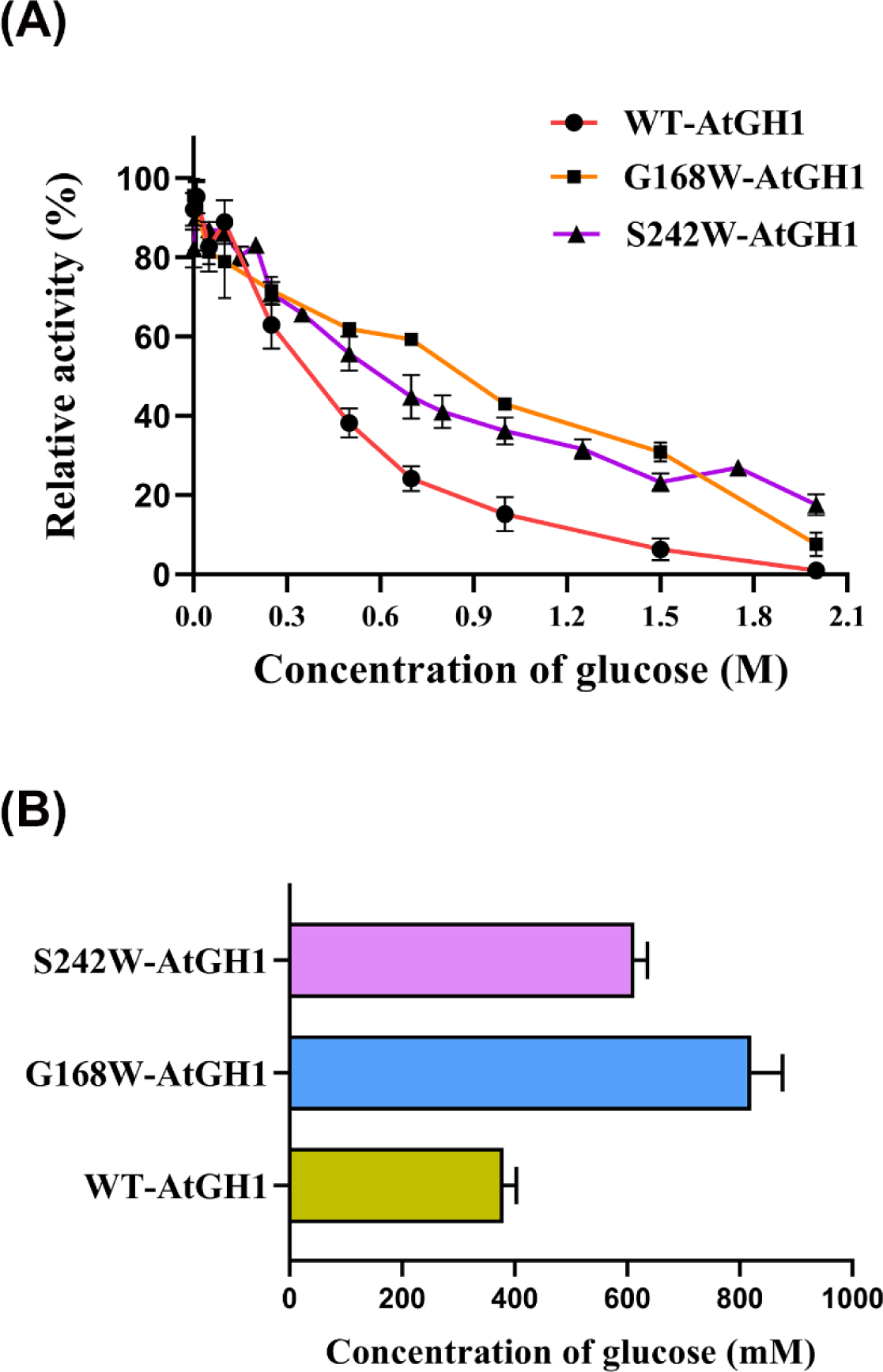
Inhibition of AtGH1 mutants by glucose. (A) Effect of glucose on the activity of WT-AtGH1, G168W-AtGH1 and S242W-AtGH1. Graph represents the relative change in the activity of WT-AtGH1 with varying concentrations of glucose. (B) Comparison of *IC*_50_ of glucose for WT-AtGH1. G168W-AtGH1 and S242W-AtGH1. The assays were done with pNPG as a substrate. Experiments were performed in triplicates (n = 3) with error bar representing ± SEM.

Gatekeeper residues at the +2 subsite lining the active site crater entry were shown to play a role in modulating the interaction of glucose molecules with the enzyme and hence affecting the glucose tolerance [37–41]. With the structural analysis of WT-AtGH1, Ser242 residue from the +2 subsite was selected for improving the glucose tolerance of WT-AtGH1. It was anticipated that an increase in the glucose tolerance of WT-AtGH1 may be possible by mutating the Ser242 to Trp. The purified S242W-AtGH1 enzyme (**Fig. S1**) showed optimum temperature for its activity at 55°C with both pNPG and cellobiose as substrates. The optimum pH for S242W-AtGH1 activity with pNPG and cellobiose as a substrate was found to be in the range of 5.5 – 6.5 (**Fig. S6**). Like G168W-AtGH1, S242W-AtGH1 was able to hydrolyse pNPG and cellobiose indicating the mutation did not lead to loss of enzyme activity (**Fig. S7**). The kinetic parameters are mentioned in table 1. S242W-AtGH1 showed an increase in the turnover number along with a threefold increase in the catalytic efficiency in comparison to WT-AtGH1 with pNPG as a substrate. Similarly, an increment in the turnover number and catalytic efficiency was observed for S242W-AtGH1 in comparison to WT-AtGH1 with its natural substrate – cellobiose, indicating that S242W mutation perturbed the kinetic properties of the WT-AtGH1 positively. Along with the enhanced kinetic parameters, the S242W-AtGH1 mutant showed improved glucose tolerance with *IC*_50_ of 612 mM glucose (**Fig. 6A and 6B**). This accounts for a ∼1.6-fold increase in the glucose tolerance level of S242W-AtGH1 as compared to that of WT-AtGH1. With the two glucose tolerant single mutants of WT-AtGH1, we decided to proceed with the double mutant – G168W-S242W-AtGH1 to achieve a cumulative effect of both these point mutations in form of superior glucose tolerance. G168W-S242W-AtGH1 mutant was generated using SDM. Surprisingly, the double mutant G168W-S242W-AtGH1 did not express in the *E. coli* expression system even after multiple trials of optimizations.

### Improvement in the kinetic properties of WT-AtGH1

While generating the S242W-AtGH1 mutant, we serendipitously obtained the S242L-AtGH1 mutant, which was confirmed by DNA sequencing. The purified S242L-AtGH1 could hydrolyse pNPG as well as cellobiose as a substrate, indicating that mutation did not lead to loss of enzyme activity. S242L-AtGH1 with pNPG as a substrate showed maximum enzyme activity at pH 5.5 with considerable activity over the pH range of 5.0 - 6.5. The mutant enzyme showed the highest activity at pH 5.5 with cellobiose as a substrate (**Fig. S6**). The mutant showed maximum activity in the temperature range of 55°C – 60°C with both pNPG and cellobiose as a substrate which is in accordance with the WT-AtGH1 enzyme (**Fig. S7**). Unlike S242W mutant, S242L showed no enhancement in the glucose tolerance level with *IC*_50_ of 393 mM (**Fig. 7A and 7B**). Kinetic parameters of S242L-AtGH1 with pNPG and cellobiose as substrates are summarized in table 1. S242L-AtGH1 mutant showed a decrease in *K*_m_ for both pNPG and cellobiose as substrates indicating the enhanced affinity of enzyme towards both the substrates. The considerable increment in the affinity of S242L-AtGH1 towards cellobiose was further complimented by ∼ 2 fold increase in the catalytic turnover (*k*_cat_) of mutant enzyme in comparison to the WT-AtGH1. The more than two fold increase in the catalytic efficiency (*k*_cat_/*K*_m_) of S242L-AtGH1 with cellobiose as compared with WT-AtGH1 highlighted the enhanced kinetic properties of S242L-AtGH1 (**Table 1**). In contrast, S242L-AtGH1 mutant showed no considerable change in the kinetic properties with pNPG as a substrate.

**Fig. 7.**
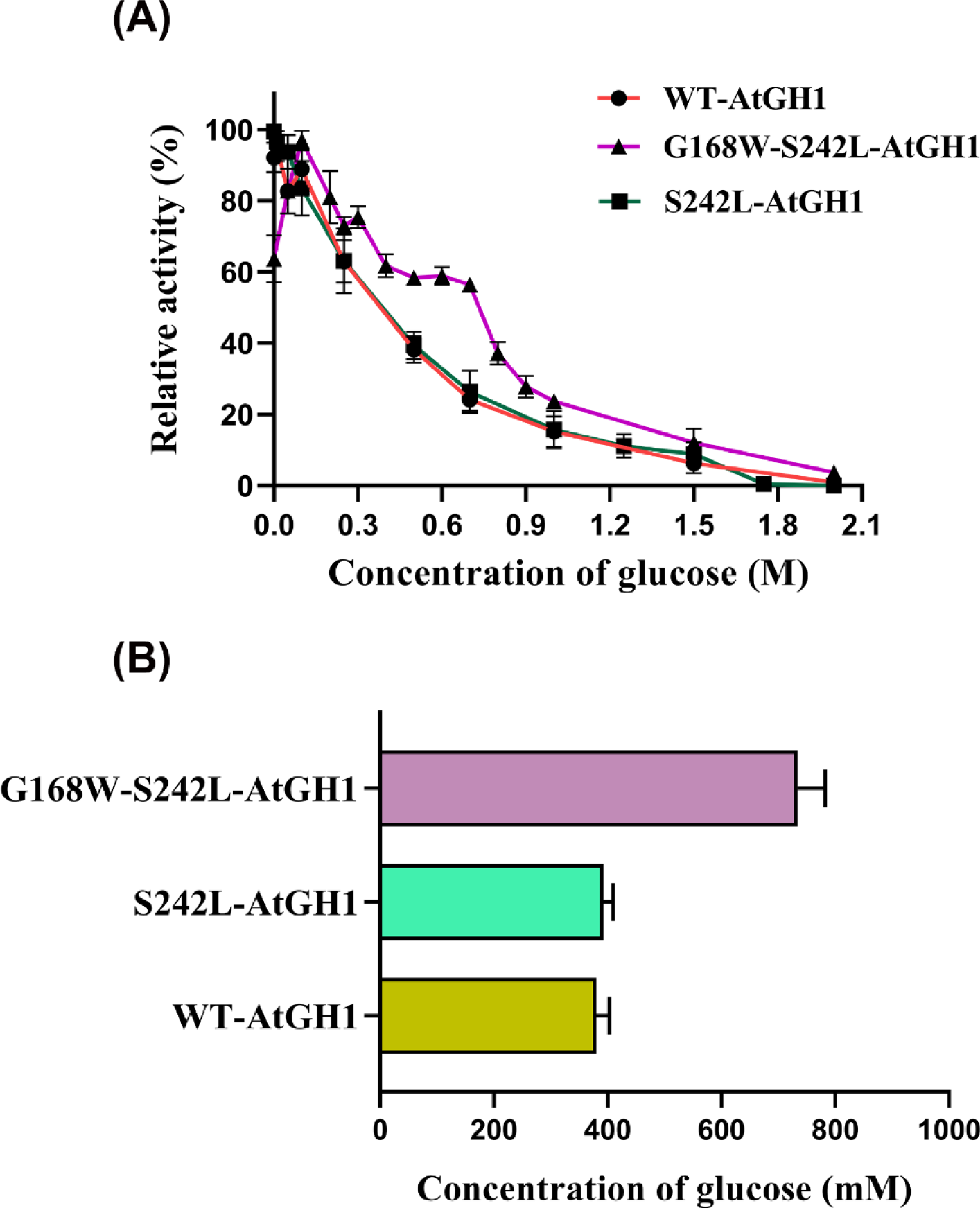
Inhibition pattern of AtGH1 mutants with improved kinetic properties by glucose. (A) Effect of glucose on the activity of WT-AtGH1, S242L-AtGH1 and G168W-S242L-AtGH1. Graph represents the relative change in the activity of WT-AtGH1 with varying concentrations of glucose. (B) Comparison of *IC*_50_ of glucose for WT-AtGH1, S242L-AtGH1 and G168W-S242L-AtGH1. The assays were done with pNPG as a substrate. Experiments were performed in triplicates (n = 3) with error bar representing ± SEM.

With the G168W mutant showing enhanced glucose tolerance and S242L mutant showing improved kinetic properties, we headed to generate a double mutant – G168W-S242L-AtGH1 that was expected to be a superior enzyme in terms of glucose tolerance as well as kinetic properties. Purified G168W-S242L-AtGH1 mutant showed activity on both – pNPG and cellobiose as a substrate (**Fig S1 and S7**). Like WT-AtGH1, G168W-S242L-AtGH1 mutant was found to show the optimum activity at 55°C in the pH range of 5.5 – 6 with pNPG and cellobiose as substrates (**Fig. S6**). The kinetic characterization of G168W-S242L-AtGH1 showed a decrease in *K*_m_ with pNPG, whereas *K*_m_ for cellobiose remained unaffected in comparison to WT-AtGH1. In contrast, the increment in the turnover number (*k*_cat_) and catalytic efficiency (*k*_cat_/*K*_m_) of G168W-S242L-AtGH1 was observed with pNPG and cellobiose (**Table 1**). Along with the improved kinetic properties, G168W-S242L-AtGH1 showed improved glucose tolerance with *IC*_50_ of 734 mM of glucose as compared to the WT-AtGH1 and S242L-AtGH1 (**Fig. 7A and 7B**).

### Thermal stability of the mutants with improved glucose tolerance and kinetic properties

For industrial usage, β-glucosidase is required to be catalytically active at an industrial operational temperature of 50°C - 60°C. Hence, we assessed the thermal stability for the mutants of WT-AtGH1.

The glucose tolerant mutants - G168W-AtGH1 and S242W-AtGH1 showed decrement in the enzyme activity by 32% and 27%, respectively, over the incubation of 48 hours at the above-mentioned conditions used for the industrial saccharification process. These values were determined by comparing with the activities of G168W-AtGH1 and S242W-AtGH1 at 0 hour of incubation. The mutant, S242L-AtGH1, with enhanced catalytic efficiency, retained 68% of the activity after incubation of 48 hours. The double mutant – G168W-S242L-AtGH1, showing enhanced glucose tolerance as well as improved catalytic efficiency, was found to be considerably active even after 48 hours of incubation at 55°C and pH 5.5, showing ∼ 74% retention of the enzyme activity (**Fig. 8**). All the mutants of WT-AtGH1 in this study exhibited high thermal stability at 55°C as well as stability in the acidic (pH 5.5) environment.

**Fig. 8.**
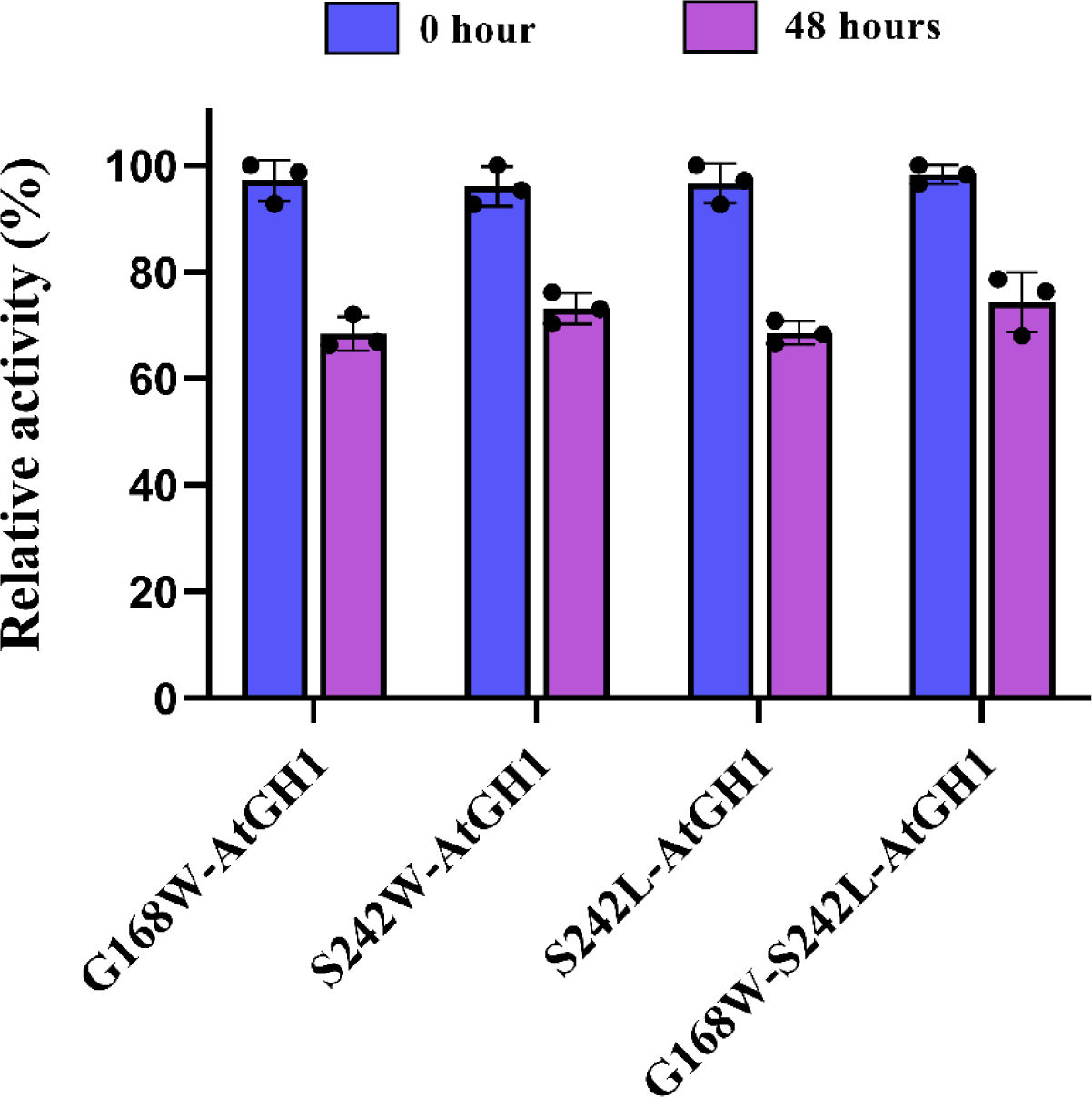
Thermal stability for mutants of WT-AtGH1 showing improved glucose tolerance and catalytic efficiency. Comparison of thermal stability of mutants at industrial operating conditions of 55 °C and pH 5.5. The bar diagram represents the relative activity of G168W-AtGH1, S242W-AtGH1, S242L-AtGH1 and G168W-S242L-AtGH1 at 0 hours (blue bar) and after 48 hours incubation at the mentioned temperature and pH (purple bar). Experiments were performed in triplicates (n = 3) with error bar representing ± SEM.

## Discussion

The saccharification of cellulosic biomass is the prime step for the industrial bioethanol production. Cellulose, being a recalcitrant polymer, requires harsh physical treatments such as steam explosion and chemical treatment methods involving acid or alkali lysis for hydrolysis to glucose. These methods include the use of corrosive chemicals, which are hazardous to personnel as well as to the environment. Hence, biochemical methods for the hydrolysis of cellulose, such as using whole-cell catalysts and cellulase enzymes, are preferred. Xiong et al. mentioned *Clostridium thermocellum* (now named as *Acetivibrio thermocellus*) as a whole cell biocatalyst for the purpose of Consolidated Bioprocessing (CBP) [42]. *Acetivibrio thermocellus* is the thermophilic anaerobic cellulolytic bacterium known to produce a cellulosomal assembly containing an extensive array of cellulases (endo-1,4-β-D-glucanases, exo-1,4-β-D-glucanases and β-glucosidases for cellulose degradation in their natural niche. Though *A. thermocellus* is utilized as a saccharifying microorganism in Consolidated Bioprocessing (CBP), the stringent restriction-modification system of the bacterium limits its genetic modifications [43] to improve its cellulose degradation ability. Hence, the focus has been moved towards isolating the cellulolytic enzymes from this bacterium and modifying those for industrial purposes.

*A. thermocellus* being a thermophilic bacterium, the extracellular enzymes produced by the bacterium such as β-glucosidases should also be able to function in high temperature environments of 50°C – 60°C. The recombinant WT-AtGH1, a β-glucosidase from *A. thermocellus* showed the optimum activity on pNPG as well as cellobiose in the temperature range of 50 °C – 60 °C. We observed distinct pH optima for WT-AtGH1 when assayed with pNPG and cellobiose as substrates. This finding has been confirmed by reproducing across three independent experimental replicates. However, the underlying mechanism for this variation in pH optima remains unclear. Kinetic analysis of the WT-AtGH1 revealed that WT-AtGH1 hydrolyses pNPG with higher catalytic efficiency than cellobiose. Though the catalytic efficiency of WT-AtGH1 for cellobiose is lower, enzyme showed prolonged activity over the period of 48 hours with least loss in the enzyme activity at high temperature of 55 °C and acidic pH of 5.5. This would aid the enzyme in efficient glucose production during the industrial saccharification, considering the operational conditions of reactors.

The main motive of this study was to develop the variants of AtGH1 with high glucose tolerance for industrial use. For that, we adopted the rational design approach of engineering this enzyme. For the rational design, we elucidated the crystal structure of WT-AtGH1. We started investigating the basis of glucose tolerance by comparing the structure of WT-AtGH1 with the structure of the glucose bound UnBGl1 - another β-glucosidase [37,38]. The UnBGl1 structure shows the presence of three glucose molecules in the active site crater at three well-defined glucose binding subsites known as −1 subsite, +1 subsite and +2 subsite at high concentrations of glucose (**Fig. S8**). This indicates that at high concentrations of product, these subsites interact with glucose molecules and, hence, may not allow cellobiose to enter the active site crater. This may lead to inhibition of enzyme activity at high concentrations of glucose. These three subsites responsible for glucose binding were also recognised in the WT-AtGH1 structure (**Fig. 5A**). Further, the sequence and structural alignment of WT-AtGH1 with other glucose tolerant glucosidases such as Bgl6, M3 mutant of Bgl6 showed the presence of hydrophobic patch with a tryptophan residue situated on the surface of the +1 subsite (**Fig. 5B**). The presence of bulky residue such as tryptophan may pose a steric hindrance to binding of a glucose molecule at +1 site. This may have led to increased glucose tolerance. The tryptophan position at the +1 subsite in Bgl6 and M3 mutant of Bgl6 was occupied by glycine in WT-AtGH1, and hence, we proposed a point mutation of G168W in WT-AtGH1 for increasing the glucose tolerance of the enzyme. This led to G168W-AtGH1 showing a two-fold increase in glucose tolerance as compared to WT-AtGH. The kinetic characterizations showed that the mutation - Gly168 to Trp led to a decrease in the turnover number and the catalytic efficiency of the enzyme with cellobiose as a substrate with no change in its affinity towards the enzyme. Cellobiose, being a natural substrate of the β-glucosidase enzyme, is a true representative of its enzyme activity. The activities of β-glucosidase with cellobiose represent the physiological relevance of the enzyme. Hence, we opine that the measurements of the activities for the characterizations of the β-glucosidase with cellobiose as a substrate should be given a preference over pNPG.

Along with the −1 subsite possessing catalytic glutamate residues and the +1 site with hydrophobic residues stabilising glucose molecule, the +2 subsite plays a crucial role in holding the glucose molecule when present in high concentrations along with dictating the entry of glucose molecule into active site crater of the enzyme. The gatekeeper residues present in the +2 site modulate glucose entry to the active site. Santos et al. introduced two mutations in the gatekeeper residues – L167W and P172L, of β-glucosidase from *Trichoderma harzianum* (ThBgl). This structure based rational approach resulted in narrowing the entry of the active site crater, enough to restrict the entry of free glucose in the active site pocket. This led to a two-fold increase in the glucose tolerance of double mutant ThBgl (*K*_i_ of 900 mM of glucose) as compared to wild-type ThBgl (*K*_i_ of 500 mM of glucose) [44]. Understanding the rationale behind mutating the gatekeeper residues, we identified Ser242 residue to introduce the mutation and enhance glucose tolerance. Following the principle of introducing bulkier residue at that position, we hypothesized the mutation of Ser242 to Trp. The modelling of S242W-AtGH1 mutant with WT-AtGH1 as a template structure showed that the diameter of the active site crater entry of S242W-AtGH1 is ∼ 16.8 Å as compared to WT-AtGH1 which is ∼ 20.5 Å (**Fig. 5C**). The narrowing of the entry is expected to hinder the entry of inhibitory glucose molecules at the +2 subsite and, hence, was expected to increase the glucose tolerance of the enzyme. The single mutation, S242W, resulted in ∼1.6 fold increase in the glucose tolerance of the enzyme in comparison to WT-AtGH1. Interestingly, the kinetic characterization demonstrated that the mutation of Ser242 in the +2 subsite led to an increment in the turnover number and catalytic efficiency of the enzyme (**Table 1**). The enhanced kinetic parameters can be seen for both substrates with a considerable improvement in case of pNPG. The positive perturbation in the kinetic parameters by a mutation in the +2 region was further confirmed by the S242L-AtGH1 mutant. With cellobiose as a substrate, S242L-AtGH1 showed enhanced affinity towards cellobiose in comparison to WT-AtGH1. This was further reflected by the ∼ 2- and 2.5-fold increase in the turnover number as well as catalytic efficiency of S242L-AtGH1, respectively (**Table 1**). As mentioned earlier, activity with cellobiose being the true representative of the catalysis by β-glucosidases, this improvement in the catalytic properties of S242L-AtGH1 is one of the important highlights of this study. This is the first time we report the improvement of catalytic parameters of β-glucosidase by a rational design approach. We strongly believe that the same amino acid position (S242 of WT-AtGH1) along with the nearby residues from other GH1 family β-glucosidases can be considered a prime position for improving their catalytic efficiencies.

Though G168W-AtGH1 and S242W-AtGH1 showed improved glucose tolerance, for their utilization in industrial saccharification processes, it is crucial to have a higher glucose tolerance, preferably exceeding 600-700 mM, considering the saccharification processes can generate glucose levels of > 800 mM [14]. With the three mutants – G168W-AtGH1, S242W-AtGH1 and S242L-AtGH1 having improved glucose tolerance and catalytic efficiency as compared to WT-AtGH1, we aimed to generate a superior enzyme by the combination of the discussed mutations. The improved glucose tolerance of G168W-AtGH1 and S242W-AtGH1 can be attributed to eliminating the inhibitory glucose molecule from +1 and +2 subsites of the active site crater, respectively. Hence, the double mutant – G168W-S242W-AtGH1 was generated and expected to have higher glucose tolerance than any of these single mutants. However, it did not express it in the *E. coli* expression system. With this, our next target was to generate another double mutant – G168W-S242L-AtGH1, which would be a combination of improved glucose tolerance and improved kinetic properties. As expected, we could observe the improvement in the kinetic properties of G168W-S242L-AtGH1 with an increase in the specific activity for cellobiose along with ∼ 1.6 fold increase in the catalytic turnover (*k*_cat_) as well as the catalytic efficiency (*k*_cat_/*K*_m_) with cellobiose as a substrate. This increment is in comparison to the WT-AtGH1. The *K*_m_ with cellobiose as a substrate for G168W-S242L-AtGH1 remained unaffected. Along with the enhanced kinetic properties, G168W-S242L-AtGH1 demonstrated ∼ 2 fold improvement in glucose tolerance level as compared to WT-AtGH1. In summary, the G168W-S242L-AtGH1 mutant was found to be the superior enzyme with its improved kinetic properties as well as glucose tolerance level. The improvement in the kinetic properties of G168W-S242L-AtGH1 with cellobiose as a substrate along with the improved glucose tolerance makes it a potential β-glucosidase candidate to be used in industrial saccharification. Along with the high glucose concentration, the operational temperature of 50°C – 60°C and operational pH of 4.8 – 5.5 are limiting factors for saccharification of cellulosic biomass in separate hydrolysis and fermentation (SHF) process of bioethanol production [22,23]. The saccharification reactors are maintained at the above-mentioned operating temperature and pH for the period of around or more than 48 hours [22,23,45]. Hence, β-glucosidase to be employed for saccharification in separate hydrolysis and fermentation (SHF) should show thermal stability over the period of 48 hours. Many of the glucose tolerant β-glucosidases show low thermostability. The half-lives of glucose tolerant β-glucosidases from *Debaromyces vanrijiae*, *Thermoanaerobacterium aotearoense* (*K*_i_ of 800 mM glucose), *Aspergillus oryzae* (*K*_i_ of 1.36 M glucose), *Candida peltata* (*K*_i_ of 800 mM glucose), a kusaya gravy metagenomic library (*IC*_50_ of > 1 M glucose), and a soil metagenomic library (*IC*_50_ of 3 M glucose) were 1, 3.3, 4, 0.5 0.2 and 1 hours at 50°C, respectively [27,46–50]. β-glucosidases from *Pyrococcus furiosus*, *Thermotoga thermarum* DSM 5069T, *Thermococcus sp*. and *Thermotoga naphthophila* RKU-10T possess both high glucose-tolerance and thermostability [51–55]. But, as these β-glucosidases are from hyperthermophilic archaea, the optimum activities of this enzyme are in the range of 80 – 95°C, which is much higher than the operating temperature range of the saccharification process, as mentioned earlier. Over these, the glucose tolerant and catalytically efficient mutants of WT-AtGH1 proved their potential in terms of retention of enzyme activities at temperature of 55°C and pH 5.5 over a period of 48 hours. As these temperature and pH conditions are maintained while operating the saccharification reactor, the mutant enzymes should be able to perform efficient hydrolysis with an average 70% retention of their enzymatic activities.

In summary, mutant enzymes under this study can be used for the saccharification of cellulose in separate hydrolysis and fermentation processes where glucose accumulates at high concentrations of > 0.6 – 0.8 M. The glucose tolerant mutants – G168W-AtGH1, S242W-AtGH1 and G168W-S242L-AtGH1 would keep on acting on the substrate – cellobiose, though a high concentration of glucose is present in the reaction environment. This would not allow cellobiose and other oligosaccharides to accumulate in the saccharification fermenter, and the upstream enzymes, such as exo- and endo-glucanase, would keep on acting on the cellulose. In addition, the thermal stability as well as stability in the acidic environment of these mutant enzymes would lower the cost of replenishing the denatured catalyst over the span of at least 48 hours for which the reactors operate. This can make the saccharification process efficient and cost-effective.

## Materials and methods

### Cloning and expression of WT-AtGH1

448 amino acid β-glucosidase protein from *Acetivibrio thermocellus*, referred to as WT-AtGH1 (GenBank - CAA42814.1), was cloned in pET28a(+) vector between *NcoI* and *XhoI* restriction sites with 6X-His tag at its C-terminus. *E. coli* DH5α competent cells were transformed with the recombinant plasmid (pET28a(+) with WT-AtGH1 gene). The recombinant clones were selected by growing transformed cells on Luria-Bertini (LB) agar containing 50 μg/ml of kanamycin. The plasmid was isolated by MiniPrep method using Exprep Plasmid SV kit (Geneall). The recombinant plasmid containing the gene for WT-AtGH1 was transformed in *E. coli* BL21 (DE3) competent cells, which were grown in 5 ml of LB containing 50 μg/ml of kanamycin at 37 °C. This actively growing culture was used to inoculate 50 ml of fresh LB broth containing 50 μg/ml of kanamycin and incubated at 37 °C with constant shaking of 150 rpm until the cells reached OD_600nm_ 0.7 - 0.8. The culture was induced with 1 mM IPTG and was further grown at 30 °C for 18 hours for the overexpression of WT-AtGH1. The cells were harvested by centrifugation at 6000g for 10 min at 4 °C. The cells were fractionated by sonication (2 sec on, 4 sec off; 40% amplitude for 10 min). The overexpression of WT-AtGH1 was confirmed by protein content analysis on SDS-PAGE [55].

### Protein Purification

AtGH1 was expressed as mentioned in above. The harvested cells were resuspended in lysis buffer - 50 mM Tris, 400 mM NaCl (pH 7.8), containing 1 mg/ml of lysozyme and 10 μg/ml of DNase at 25 °C with constant shaking for 30 min followed by sonication (2 sec on, 4 sec off; 40% amplitude for 10 min). The cell lysate was centrifuged at 16000g for 30 min at 4°C and the supernatant collected was filtered through 0.45 μm PVDF filter. The filtrate was loaded on Ni-NTA affinity chromatography column pre-equilibrated with an equilibration buffer (50 mM Tris, 400 mM NaCl, pH 7.8). After loading, the column was thoroughly washed with a buffer containing 50 mM Tris, 400 mM NaCl and 50 mM imidazole to remove non-specifically bound protein. Further, WT-AtGH1 was eluted using 50 mM Tris, 400 mM NaCl (pH 7.8) containing increasing concentrations of imidazole (100 mM, 175 mM and 250 mM). Fractions containing WT-AtGH1 were pooled and concentrated using Amicon ultrafiltration concentrator (∼ 10kDa cut off) followed by the gel filtration chromatography purification. The sephacryl S200 10/300 gel filtration column pre-equilibrated with buffer pH 7.0 (50 mM Tris, 50 mM NaCl) was loaded with the concentrate. The elution fractions containing WT-AtGH1 were collected, pooled and stored at 4°C. After each purification step, the purity and activity of WT-AtGH1 were assessed by SDS-PAGE and enzyme activity studies (as mentioned in the later sections). The concentration of purified protein was quantified by the Bradford method of protein estimation [56].

### Biochemical and kinetic characterization of WT-AtGH1

#### Enzyme activity assay

For the functional characterization of WT-AtGH1, *p*-nitrophenyl-β -D-glucoside (pNPG) (Merck) and cellobiose (Sigma-Aldrich) were used as substrates. The assays were performed with 50 μl of 40 mM pNPG stock diluted in 440 μl of 50 mM sodium acetate buffer, pH 5.5. Reaction mixtures were preincubated at 55 °C for 15 min. Further, 10 μl of 0.05 mg/ml enzyme was added to the above reaction mixture. The reaction was incubated at 55 °C for 10 min. The reaction was terminated by adding 500 μl of 0.2 M sodium carbonate. The released product *p*-nitrophenol (pNP) was estimated spectrophotometrically by measuring absorbance at 405 nm. One enzyme unit corresponds to the amount of enzyme required to hydrolyse 1 µmole of pNPG per minute under the mentioned assay conditions. The natural substrate for β-glucosidase - cellobiose was prepared in 50 mM sodium acetate buffer, pH 5.5. The assays were performed in 0.2 ml microfuge tubes with 95 μl of 350 mM cellobiose stock. 5 μl of 1 mg/ml enzyme was added to the same 0.2 ml microfuge tube and the reaction was incubated at 55 °C for 30 min. Reactions were terminated by keeping the reaction mixture in the thermocycler at 95°C for 15 min. Reaction mixtures were cooled and 2 μl of an aliquot from the reaction mixture was mixed with 200 μl of GOD-POD reagent (Eco Pak Glucose 500, Accurex Biomedicals Pvt. Ltd.) in 96 well microtitre plate. This was incubated at 37°C for 15 min. The pink colour developed was estimated spectrophotometrically in the microplate reader (Vantastar, BMG Labtech) at 505 nm for calculating enzyme activity. One enzyme unit corresponds to the amount of enzyme required to hydrolyse 1 µmole of cellobiose into glucose per minute under mentioned assay conditions. The enzyme activities are calculated as specific activity of the enzyme with pNPG or cellobiose as substrates and expressed as µmole/min/mg.

#### Determination of optimum temperature and pH

The optimum temperature for the activity of WT-AtGH1 was determined by carrying out the reactions in 50 mM sodium acetate buffer, pH 5.5 at varying temperatures - 30°C, 40°C, 50°C, 55°C, 60°C, 70°C, 80°C in. The optimum pH for the activity of WT-AtGH1 was estimated by carrying out reactions at 55°C and varying pH - pH 4.0, 4.5, 5.0, 5.5 (50 mM sodium acetate), pH 6.0, 6.5, 7.0, 7.5, 8.0 (50 mM sodium phosphate buffer). Reactions were carried out as mentioned above.

#### Enzyme kinetics studies

The kinetic characterization of the WT-AtGH1 was done using two substrates – pNPG and cellobiose. For pNPG, enzymatic reactions were carried out in 50 mM sodium acetate buffer pH 5.5 with varying concentrations of pNPG (100 μM to 10 mM), keeping enzyme concentration (10 µl of 0.05 mg/ml) the same. With cellobiose as substrate, reaction mixtures constituted 5 µl of 1 mg/ml enzyme along with various concentrations of cellobiose (3.5 mM to 332.5 mM). The assays were performed as mentioned previously. The specific activities for each of the substrate concentrations were calculated and plotted against the concentration of substrate. The kinetic parameters – *K*_m_ and *V*_max_ were determined by fitting the data to the appropriate kinetic model using GraphPad Prism 8.4.2.

#### Effect of metal ions

The effect of metal ions on the activity of WT-AtGH1 was assessed by carrying out reactions with pNPG in the presence of varying concentrations (1mM, 5 mM, 10 mM and 50 mM) of metal ions – Mg^2+^, Na^+^, Ca^2+^, Fe^2+^, Hg^2+^, Cu^2+^, Zn^2+^, Co^2+^, K^+^ and Mn^2+^. The enzyme was preincubated with corresponding concentrations of metal ions for 30 min before assays. The reactions and assays were carried out with pNPG as mentioned above.

#### Determination of glucose tolerance

The inhibition pattern of WT-AtGH1 with glucose as an inhibitor was determined using pNPG as a substrate. The enzyme assays were performed with reaction mixtures containing varying concentrations of glucose (10 mM – 2 M). Each reaction mixture of 500 μl constituted a fixed concentration of glucose with 2 mM pNPG and 10 µl of 0.05 mg/ml enzyme in 50 mM sodium acetate buffer pH 5.5. The saturation kinetic studies of WT-AtGH1 with pNPG were carried out with three different concentrations of glucose (40 mM, 120 mM and 200 mM). Inhibition pattern was determined from Lineweaver-Burk plot and *K*_i_ was calculated from Dixon plot.

#### Assessment of the thermal stability

The thermal stability of WT-AtGH1 was assessed by incubating the enzyme in 50 mM sodium acetate buffer, pH 5.5 in a water bath at 55 °C. Enzyme assays were performed using 350 mM cellobiose as substrate. WT-AtGH1 (1 mg/ml) was incubated at 55 °C in 50 mM sodium acetate buffer pH 5.5 for a duration of 48 hours. 5 µl of the incubated enzyme was taken for the enzyme assay. The specific activity of WT-AtGH1 at 0 hour and at 48 hours was calculated and compared to measure its thermal stability.

### Structure elucidation of WT-AtGH1

#### Protein Crystallization

Crystallization screening of WT-AtGH1 was carried out with concentrated (∼ 10mg/ml) protein in a 96-well polystyrene plate with a sitting drop vapour diffusion method. The screening was set up with commercial screens - JCSG+, PEG Rx, PEG/Ion, Index and PEG suite (Molecular Dimensions) by mixing 0.2μl of AtGH1 and 0.2 μl mother liquor equilibrated against 50 μl of the same reservoir solution with the help of crystallization robot, Phoenix (Art Robbins Instruments) at Protein Crystallography Facility, IIT Bombay. The crystallization plates were incubated at 22°C. Initial crystals were obtained in 0.1 M HEPES (pH 7.5) and 70% MPD. Further optimization was carried out in 24 well poly-styrene plates by hanging drop vapour diffusion method at 22°C. An optimized condition with 0.1 M HEPES (pH 7.5) and 55 % MPD showed the appearance of better-quality crystal after 8 days of incubation at 22°C.

#### Diffraction data collection, structure solution and refinement

The X-ray diffraction data collection was performed at 100 K with mother liquor itself as a cryoprotectant under a stream of liquid nitrogen cryo-conditions. A crystal was picked from the crystallization well using a cryo-loop and was flash-cooled into the stream of liquid nitrogen. The data set was collected with 0.5° rotation per frame at a wavelength of 1.5418 Å by Cu - *K*α X-ray radiation generated by a Rigaku Micromax 007HF diffractometer equipped with R-AXIS IV++ image plate detectorat Protein Crystallography Facility, IIT Bombay. Data set collected in the form of frames were indexed and integrated using XDS program [57]. These integrated intensities were converted to structure factor in mtz format using XDSCONV. Hence, phases were calculated using the structure of β-glucosidase from *C. cellulovorans* (PDB entry 3AHX, sequence identity 56%). The molecular replacement method of PHASER was used for calculating phases using CCP4 [58,59] and the online server BALBES [60]. Further structure refinement was performed using REFMAC5 from CCP4 and phenix.refine from PHENIX [61–63]. After each cycle of refinement, the model was rebuilt using COOT [64,65]. Solvent molecules were added in high peak electron densities in σ_A_ weighted *F*_o_ – *F*_c_ map. Improvements in model rebuilding were monitored by an increase in figures of merit with a decrease in *R*_free_. Final cell parameters, data processing and refinement statistics are summarised in **Table 2**. Structural alignments were done using the SSM tool of COOT, and figures for the structure were prepared using PyMOL [66] and UCSF ChimeraX [67,68].

### Site directed mutagenesis, expression and purification of WT-AtGH1 mutants

The site directed mutagenesis of WT-AtGH1 was performed by PCR for generation of single mutants - G168W, S242W, S242L and double mutants - G168W-S242W and G168W-S242L. pET28a(+) vector harbouring the WT-AtGH1 gene was used as a template for generating these mutants. The primers used for the mutagenesis are listed in **table S1**. The mutations introduced were confirmed by DNA sequencing. The expression studies of the AtGH1 mutants were performed using the same procedure followed for WT-AtGH1. The above-mentioned WT-AtGH1 mutants (except G168W-S242W which did not express) were purified to homogeneity by Ni-NTA affinity chromatography followed by gel filtration chromatography. After each step of purification, the purity and activity of mutants were assessed by SDS-PAGE, and enzyme activity studies were carried out, as mentioned earlier.

### Biochemical and kinetic characterization of WT-AtGH1 mutants

The biochemical characterizations of the above-mentioned mutants of WT-AtGH1, including the determination of temperature and pH optimum for the activity, extent of glucose inhibition in terms of *IC*_50_ calculations, the kinetic characterizations and thermal stability assays, were performed as mentioned for WT-AtGH1.

## Acknowledgements

Funding for the above study has been provided by the DBT PAN-IIT Centre for Bioenergy; Grant from the Department of Biotechnology (DBT), Government of India, Grant number - BT/PR41982/PBD/26/822/2021. We acknowledge Prof. Arun Goyal (Indian Institute of Technology Guwahati) for generously providing us with the plasmid construct containing the AtGH1 gene. We acknowledge the support from the Protein Crystallography Facility at IIT Bombay for protein crystallization and X-ray diffraction data collection. CK acknowledges IIT Bombay for providing Ph.D. fellowship.

## Author contributions

CK: Conceptualization, Protein purifications, biochemical and kinetics assays, data collection and structure solution, validation, data analysis, writing original draft, reviewing and editing the draft

AR: Protein purifications, biochemical and kinetics assays, validation, data analysis, reviewing and editing the draft

PB: Conceptualization, data collection and structure solution, validation, data analysis, writing original draft, reviewing and editing the draft, resources, funding acquisition, project review and administration, supervision

## Data availability statement

This article contains supporting information. Structure factors and atomic coordinates have been submitted to Protein Data Bank (PDB) under PDB ID – 9IQB. Most of the data are contained within the article itself. Any other data not included in the article can be made available by the corresponding author upon reasonable request.

