## Supplementary information for "Rational design paving the way for improving glucose tolerance and catalytic properties of a β-glucosidase from *Acetivibrio thermocellus*"

**Running title:** Rational engineering of β-glucosidase from *Acetivibrio thermocellus*

**Keywords:** β-glucosidase, GH1 family, crystal structure, cellobiose, glucose tolerance, thermal stability, catalytic efficiency

### Supplementary materials and methods

#### Circular dichroism analysis of WT-AtGH1

Far-UV CD spectra of WT-AtGH1 were recorded between the wavelength range of 195 nm – 260 nm in a cuvette of path length 0.1 cm using a spectropolarimeter (J-1500, Jasco). 5 μM of WT-AtGH1 made in 50 mM sodium acetate buffer of pH 5.5. WT-AtGH1 was incubated at 55C for 48 hours and the CD spectrums were recorded at 0 hour and 48 hours of incubation. Parameters set for recording CD spectrum - sensitivity of 100 millidegrees, scan speed of 100 nm/min, response time of 2 seconds with three accumulations.

| Primers used for SDM | Primer sequence |
|---|---|
| G168W-AtGH1-FP | 5′ GAACCC**TGG**GTTGTTTCTTTGCTTGGCCACTTTTTAGG 3′ |
| G168W-AtGH1-RP | 5′ GAAACAAC**CCA**GGGTTCATTGTGAGTAAACCATATTGG 3′ |
| S242W-AtGH1-FP | 5′ GCTGAGGATATTGAAGCAGCGGAATTG**TGG**TTTTCTCTGG 3′ |
| S242W-AtGH1-RP | 5′ GATACCACCTTCCCGCCAGAGAAAA**CCA**CAATTCCGCTGC 3′ |
| S242L-AtGH1-FP | 5′ GCTGAGGATATTGAAGCAGCGGAATTG**CTG**TTTTCTCTGG 3′ |
| S242L-AtGH1-RP | 5′ GATACCACCTTCCCGCCAGAGAAAA**CAG**CAATTCCGCTGC 3′ |

**Table S1.** Primers used for site directed mutagenesis for the generation of mutants – G168W-AtGH1, S242W-AtGH1, S242L-AtGH1 and G168W-S242L-AtGH1.

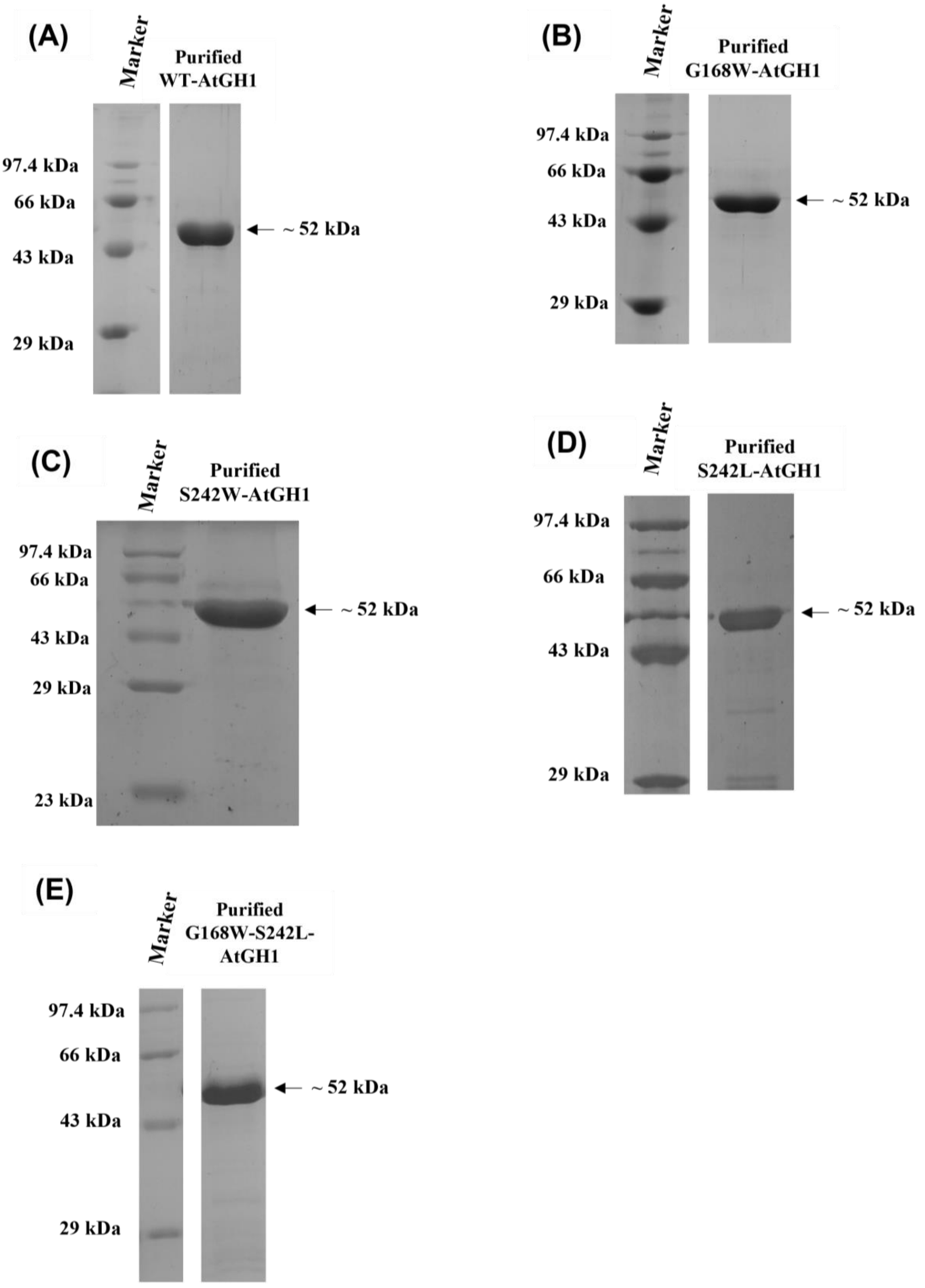


**Figure S1.** SDS-PAGE gel for the (A) purified WT-AtGH1, (B) purified G168W-AtGH1, (C) purified S242W-AtGH1, (D) purified S242L-AtGH1, (E) purified G168W-S242L-AtGH1 after the final purification step of gel filtration chromatography.

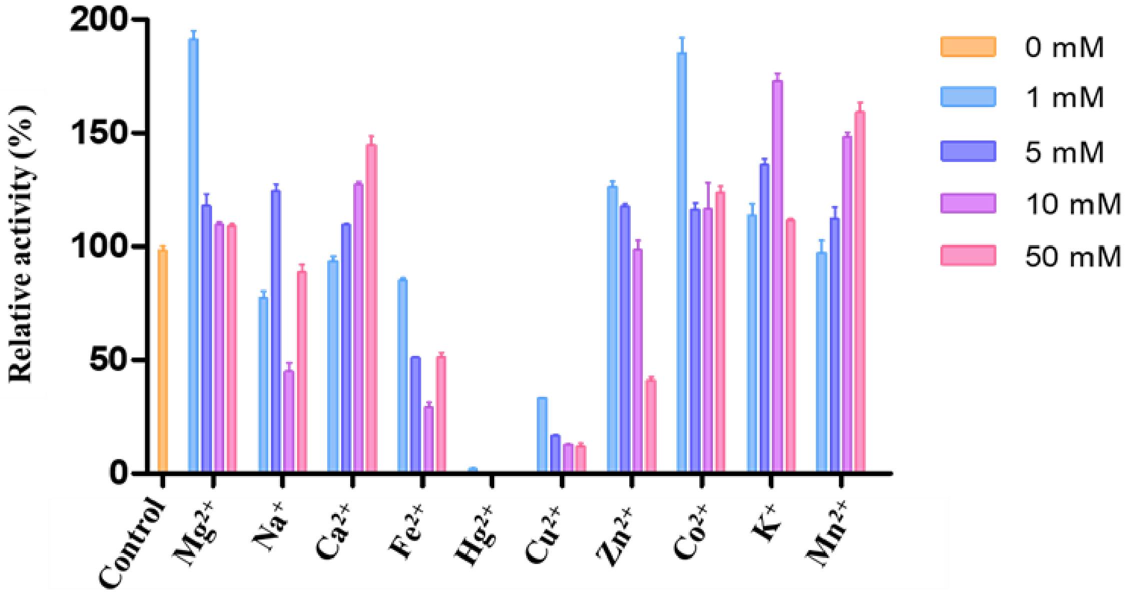


**Figure S2.** Effect of various metal ions on the activity of WT-AtGH1. The graph shows the relative activity in presence of four different concentrations of metal ions. The assay for the enzyme was performed with pNPG as a substrate. Experiments were performed in triplicates (n = 3) with error bar representing ± SEM.

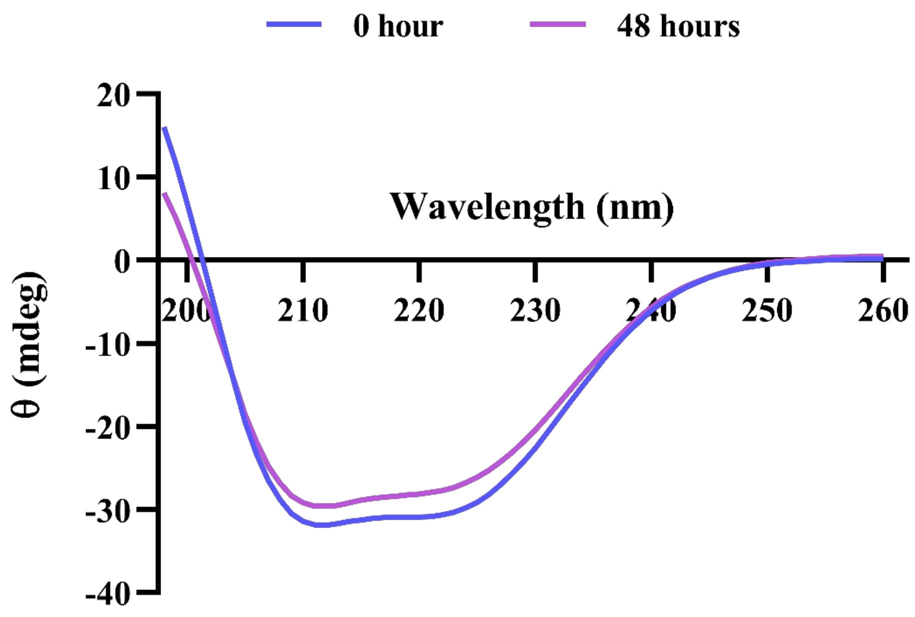


**Figure S3.** The circular dichroism (CD) spectrum of WT-AtGH1. The graph represents ellipticity recorded over a specified wavelength range. WT-AtGH1 was incubated at 55°C for 48 hours. CD spectrum was recorded at 0 hour (blue curve) and after 48 hours (purple curve) of incubation at mentioned conditions.

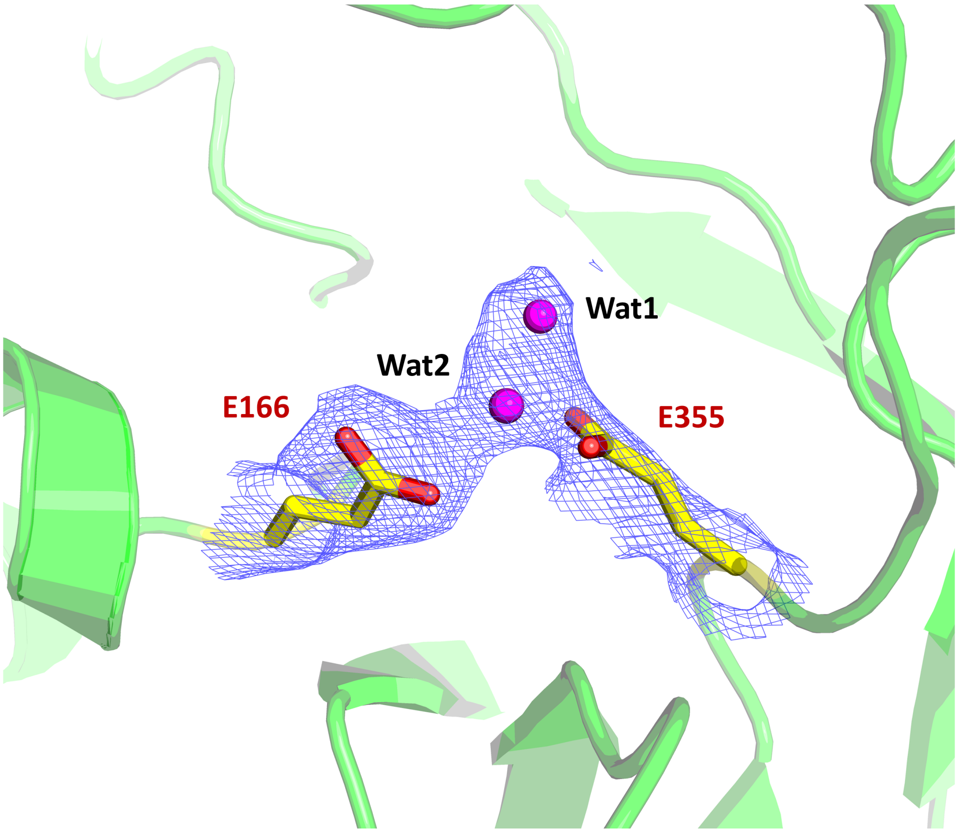


**Figure S4.** $2F_o$-$F_c$ map (contour level 1σ) in blue mesh is shown around two water molecules – Wat1 and Wat2 (magenta spheres) and closely associated with catalytic glutamates – E166 and E355. The residues are as represented as yellow sticks.

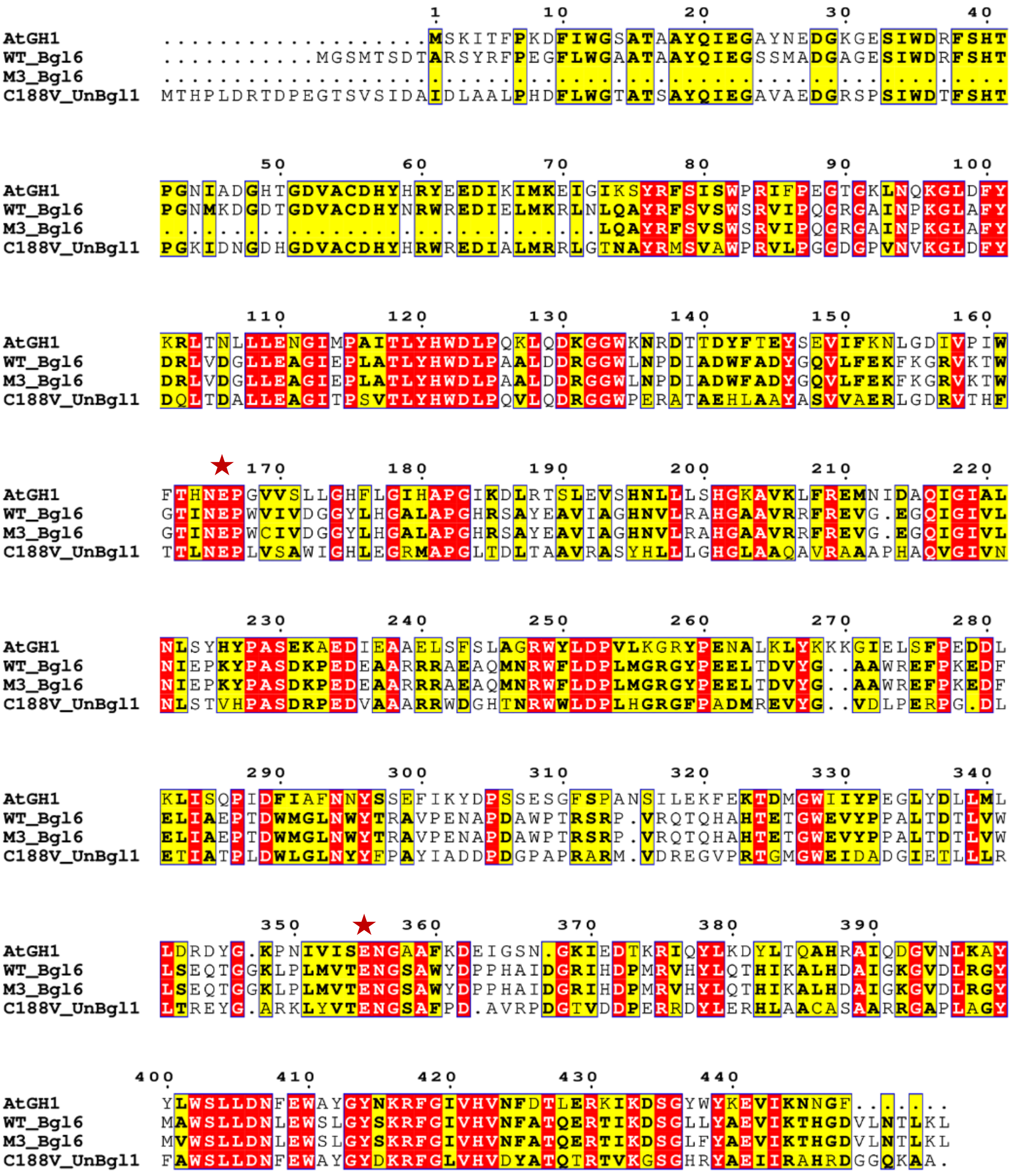


**Figure S5.** Multiple sequence alignment of WT-AtGH1 with Bgl6, M3 mutant of Bgl6 and C188V mutant of UnBGl1. Red stars show two catalytic glutamates – E166 and E355 from conserved motifs – NEP and ENG, in all mentioned β-glucosidases. Sequences retrieved – WT_Bgl6 (PDB entry 5GNY), M3_Bgl6 (PDB entry 5GNZ), C188V_UnBGl1 (NCBI accession number - AFU63315.1), AtGH1 (NCBI accession number - CAA42814.1); sequence alignment is generated from ESPript 3.0 server [1].

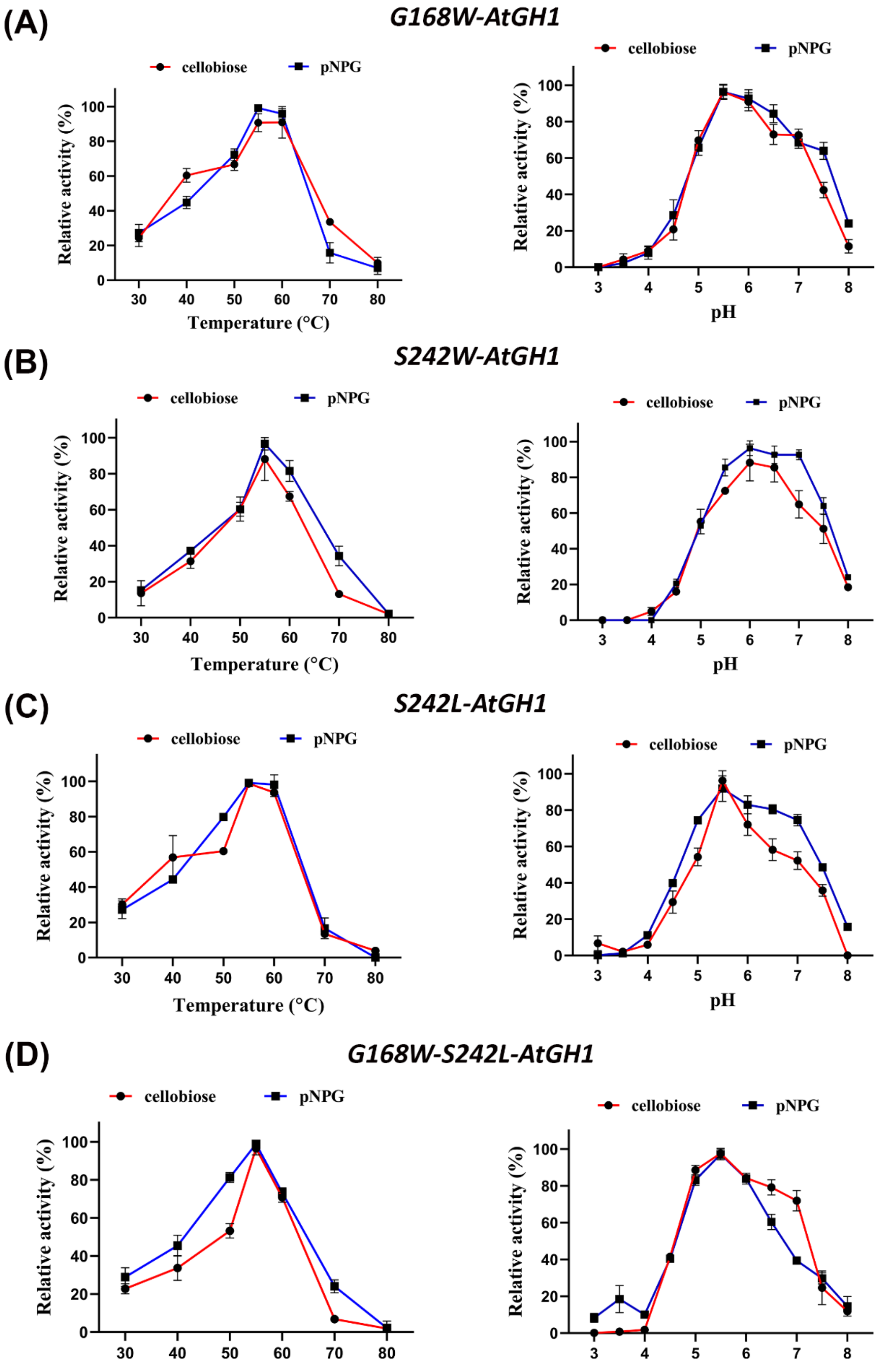


**Figure S6.** Effect of temperature (left panels) and the effect of pH (right panel) on the activities of (A) G168W-AtGH1, (B) S242W-AtGH1, (C) S242L-AtGH1 and (D) G168W-S242L-

AtGH1. The assays were done with cellobiose (red curve) and pNPG (blue curve) as substrates. All experiments were performed in triplicates (n = 3) with error bar representing ± SEM.

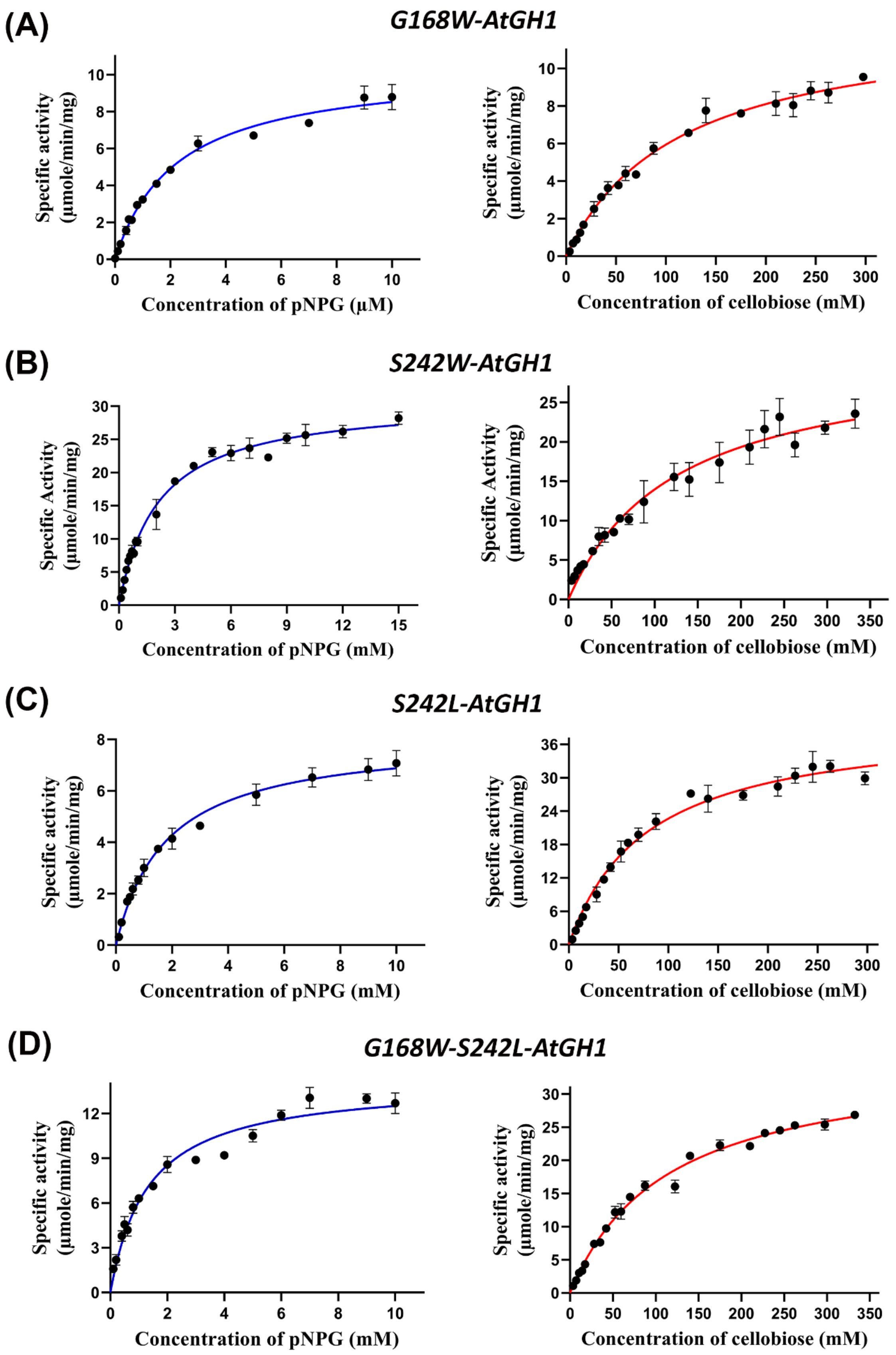
(A)
G168W-AtGH1
Specific activity (µmole/min/mg)
Concentration of pNPG (µM)
Specific activity (µmole/min/mg)
Concentration of cellobiose (mM)
(B)
S242W-AtGH1
Specific Activity (µmole/min/mg)
Concentration of pNPG (mM)
Specific Activity (µmole/min/mg)
Concentration of cellobiose (mM)
(C)
S242L-AtGH1
Specific activity (µmole/min/mg)
Concentration of pNPG (mM)
Specific activity (µmole/min/mg)
Concentration of cellobiose (mM)
(D)
G168W-S242L-AtGH1
Specific activity (µmole/min/mg)
Concentration of pNPG (mM)
Specific activity (µmole/min/mg)
Concentration of cellobiose (mM)

**Figure S7.** Enzyme kinetics of (A) G168W-AtGH1, (B) S242W-AtGH1, (C) S242L-AtGH1 and (D) G168W-S242L-AtGH1. The assays were done with pNPG (blue curve) and cellobiose (red curve) as substrates. All experiments were performed in triplicates (n = 3) with error bar representing ± SEM.

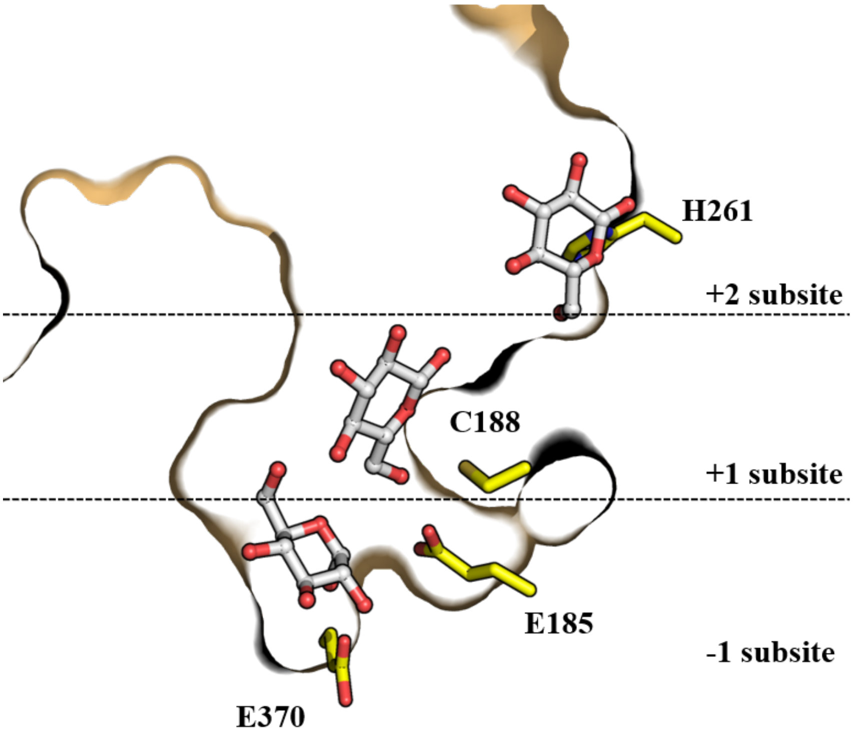


**Figure S8.** Cross section of the active site crater of wild type UnBGl1 showing the presence of three glucose molecules (represented as balls and sticks with gray) at conserved -1, +1, and +2 subsites. Catalytic residues of UnBGl1 – E185 and E370 are shown in ball and stick along with C188 and H261 forming a part of +1 and +2 subsites [2,3].
